# Effects of input image size on the accuracy of fish identification using deep learning

**DOI:** 10.1101/2024.03.01.582886

**Authors:** Yuka Iwahara, Yasutoki Shibata, Masahiro Manano, Tomoya Nishino, Ryosuke Kariya, Hiroki Yaemori

**Affiliations:** Fisheries Resources Institute, National Research and Development Agency, Japan Fisheries Research and Education Agency, Yokohama, Kanagawa, Japan; Computermind Corp., Nishi-Shinjyuku, Tokyo, Japan

**Keywords:** Mask R-CNN, Species classification, Input image size, Set-net, Stock assessment

## Abstract

The length composition of catches by species is important for stock assessment. However, length measurement is performed manually, jeopardizing the future of continuous measurement because of likely labor shortages. We focused on applying deep learning to estimate length composition by species from images of fish caught for sustainable management. In this study, input image sizes were varied to evaluate the effect of input image size on detection and classification accuracy, as a method for improving the accuracy. The images (43,226 fish of 85 classes) were captured on conveyor belts to sort set-net catches. Fish detection and classification were performed using Mask R-CNN. The effect of input image size on accuracy was examined using three image sizes of 1333×888, 2000×1333, and 2666×1777 pixels, achieving an mAP50-95 of 0.580 or higher. The accuracy improved with increasing image size, attaining a maximum improvement of 4.3% compared to the smallest size. However, increasing the image size too far from the default size may not improve the accuracy of models with fine-tuning. Improvements in accuracy were primarily observed for the species with low accuracy at the smallest image size. Increasing image size would be a useful and simple way to improve accuracy for these species.

## Introduction

The appropriate utilization of fisheries stock is essential for their sustainable exploitation. Therefore, stock assessments should be conducted to determine whether a catch is suitable for the current stock status. More accurate stock assessments require not only catch amounts by fish species but also fish length composition (e.g. Arreguin-Sanchez, 1996; Ono *et al*., 2015; Wetzel & Punt, 2011). Length information is collected by manually measuring the fish sampled from research vessels or catches in fishing ports (Øvredal and Totland, 2002; Palmer *et al*., 2022).

The number of fish species and the sample size that can be manually measured depends on the number of people measuring. Additionally, these measurements require skilled researchers to identify the fish species and examine their lengths. Moreover, these measurements must be performed continuously because stock assessments are regularly conducted. In recent years, some countries have experienced a shortage of workers for reasons such as human population decline and aging (Challenger, 2003; Ducanes and Abella, 2008). The shortage of skilled researchers is a serious problem, leading to increased uncertainty in stock assessments. Therefore, the development of a method that does not depend on the number of researchers is important for stock assessment and sustainable exploitation of fish stocks.

In recent years, deep learning approaches using images have been garnering interest as methods for replacing manual methods of classification, inspection, and size estimation in many fields such as agriculture, industry, and medicine (e.g. Fuentes *et al*., 2017; Shen *et al*., 2019; Tang *et al*., 2020; Blok *et al*., 2021). Deep learning has also been used in fisheries science to examine the amount of discarded fish that have not landed or catches of juvenile fish that are difficult to identify by species in longline, purse seine, and trawl fisheries in Europe and the U.S. (e.g. French *et al*., 2020; Mei *et al*., 2021; Lekunberri *et al*., 2022).

However, few studies have used deep learning to estimate fish length for stock assessment (e.g. Álvarez-Ellacuría *et al*., 2020; Palmer *et al*., 2022). Shibata *et al*. (2024) used deep learning approaches to collect fish length information as a parameter for stock assessment, estimating the length with an accuracy of ±3%. They took photographs of the species after the catches had been manually identified and did not conduct species identification using deep learning.

In previous studies, fish species identification has been performed using images taken onboard, as the purpose of these studies was electronic monitoring of fishery activities, such as discards and the amount of catches. In most fishing gear, there are catches of multiple fish species; therefore, fish species identification is crucial when using images of catches for stock assessment. Nets of purse seine or trawls usually capture large numbers of fish and multiple species simultaneously (French *et al*., 2020; Lekunberri *et al*., 2022). Images of the fish caught by these fishing gears were captured using cameras installed on onboard conveyors for sorting fish or onboard conveyors specifically for image capture (French *et al*., 2020; van Essen *et al*., 2021; Lekunberri *et al*., 2022). However, conveyors specifically for imaging require spaces on board. Even if only a camera were installed, fishermen working or sorting on small ships would block to capture the catches. Therefore, onboard methods applied in previous studies to large fishing vessels, are difficult to apply to small fishing vessels. There are fewer large fishing vessels in Asia than in Europe or the U.S. (FAO, https://www.fao.org/3/cc0461en/online/sofia/2022/fishing-fleet.html, 3/1/2024), making it difficult to use onboard cameras, as in previous studies.

Length information for stock assessment is collected not only from vessels but also from fishing ports (Mace *et al*., 2001). Therefore, fishing ports are potential locations for capturing images of catches. In particular, set-nets catch many fish species (e.g. Training Department of Southeast Fisheries Development Center, 2008; Lu and Lee, 2014; Ministry of Agriculture, 2023), so those catches are generally sorted at fishing ports. Information on numerous fish species can be obtained by capturing images from set-net catches. This is considered an advantage when collecting information for stock assessment.

In general, the larger the input image size, the better the classification accuracy (Huang *et al*., 2018). Previous studies in fields other than fisheries have also demonstrated improved accuracy in discriminating the shape of coffee beans and classifying leaf types (Camgözlü and Kutlu, 2020; Gope and Fukai, 2022). Increasing the image size would be useful for fish species classification because the characteristics used for identification are often very small. In addition, smaller objects tend to be missed more often in object detection (Kisantal *et al*., 2019). The camera fixed on a conveyor belt captures images of various-sized fish from a fixed distance; therefore, the smaller the fish, the smaller the fish size in the image. Therefore, verifying the extent to which detection and classification accuracy can be improved by increasing the size of the input images is important. However, no attempts have been made to change the image size in fish detection and classification. Set-nets, with a wide variety of species in their catches, are suitable for verifying the species that can be more accurately identified when image size is increased.

Therefore, our objective was to evaluate the effect of increasing the image size on the accuracy of fish detection and species classification. Our study is expected to contribute to improving the accuracy of object detection and classification of fish images, thereby contributing to the estimation of fish length information for stock assessment using catch images.

## Materials & Methods

### Collection of fish images

Fish images were obtained from set-net catches sorted using a conveyor belt at the Odawara fishing port in Kanagawa Prefecture, Japan, from August 2020 to October 2021 (Figure 1). A camera (DSC-RX0M2, Sony) was fixed 70 cm from the surface of the conveyor belt, and images were captured automatically once per second. Images were obtained between approximately 4:00 - 7:00 am, when the catch landed. The image size was 4800×3200. The shutter speed and exposure compensation were manually adjusted before capturing the images to avoid blurring or hiding the color pattern of the fish due to darkness or reflections of light. The shutter speed was set to a range between 1/320–1/100 of a second. LED lights (Sanwa Supply Inc., 200-DG019) and polarizing filters were used as required.

**Fig. 1.**
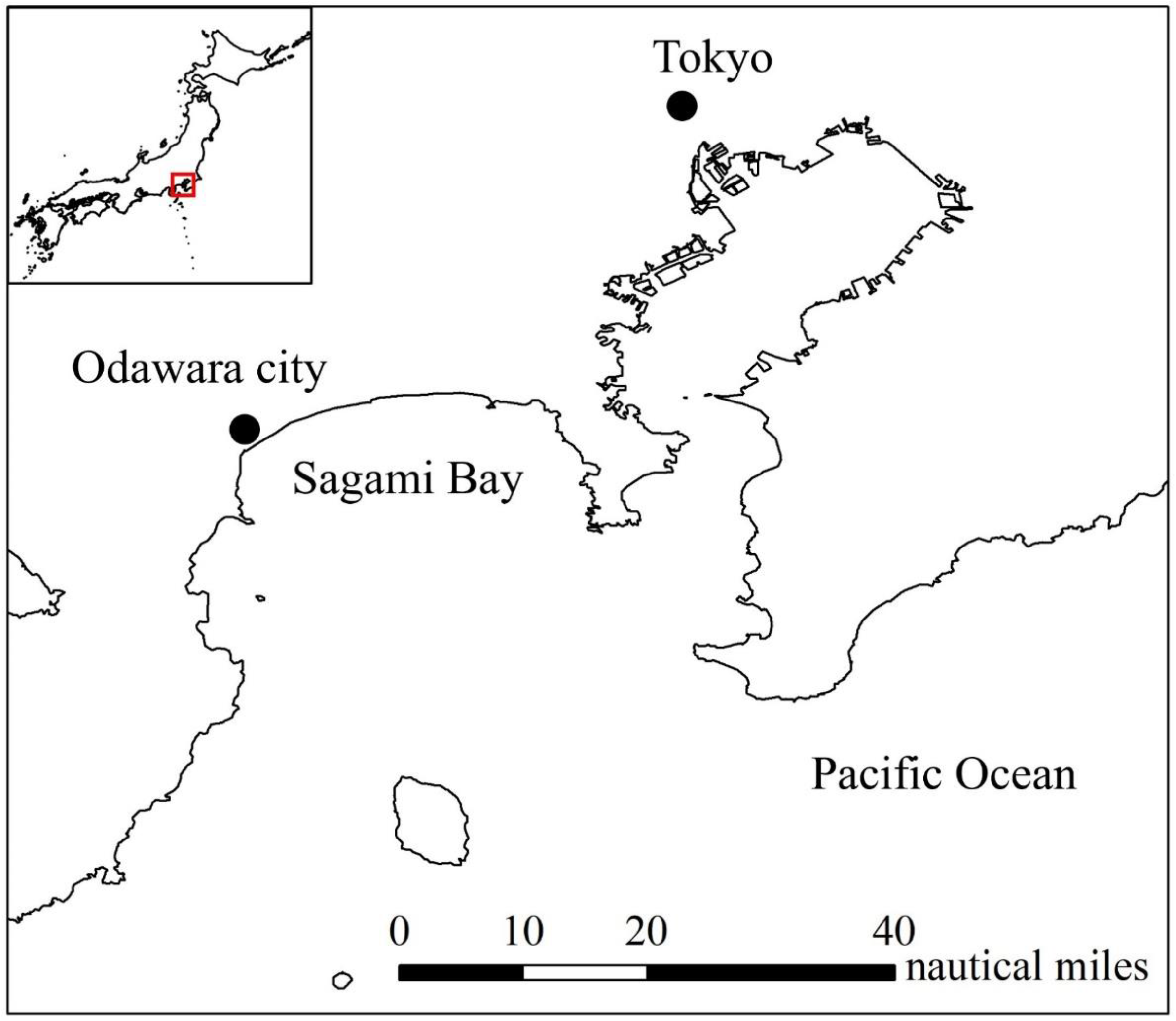
Position of Odawara city.

### Annotation and classes

The images were annotated using instance segmentation. Each fish was labeled at the species level whenever possible. When species could not be identified, they were grouped together based on their similar appearance at the family, genus, or higher taxonomic levels (Supplemental Table 1). Because juvenile and adult chicken grunts (*Parapristipoma trilineatum*) have different body color patterns, they were placed in separate classes (chicken grunt juvenile and chicken grunt). The total number of data consisted of 3,909 images from 85 classes and 43,226 individuals (Table 1). To avoid overfitting, fish that consistently appeared in the same positions under the conveyor belt in multiple images, were excluded from the analysis. The excluded fish were filled with simple square black polygons so that they could be treated as background in the training and not as fish.

**Table 1.** List of fish classes and number of data.

| Anotation name | Train | Validation | Test | Total | Anotation name | Train | Validation | Test | Total |
| --- | --- | --- | --- | --- | --- | --- | --- | --- | --- |
| Red stingray | 16 | 1 |  | 17 | Rusty jobfish | 2 |  |  | 2 |
| Lesser moray eel | 15 | 3 | 2 | 20 | Double-lined fusilier | 3 | 1 | 1 | 5 |
| Daggertooth pike conger | 40 | 5 | 17 | 62 | Chicken grunt | 121 | 10 | 8 | 139 |
| Bali sardinella | 10 | 1 |  | 11 | Chicken grunt juvenile | 1024 | 128 | 79 | 1231 |
| Japanese sardine | 238 | 32 | 21 | 291 | Crimson sea-bream | 253 | 35 | 36 | 324 |
| Round herring | 767 | 174 | 158 | 1099 | Goldlined seabream | 5 |  | 3 | 8 |
| Japanese anchovy | 48 | 6 | 7 | 61 | Grey large-eye bream | 6 | 2 | 1 | 9 |
| Barbel eel | 54 | 9 | 5 | 68 | Bensasi goatfish | 10 | 3 | 1 | 14 |
| Brushtooth lizardfish | 63 | 4 | 5 | 72 | Bluestriped angelfish | 3 |  |  | 3 |
| "Eso" | 5 | 1 |  | 6 | Yellowstriped butterflyfish | 3 | 1 | 1 | 5 |
| "Iwashi" | 35 | 1 | 6 | 42 | Stripey | 15 |  | 4 | 19 |
| Myctophidae | 50 | 7 | 10 | 67 | Dotted gizzard shad | 56 |  | 6 | 62 |
| "Tobiuo" | 74 | 8 | 4 | 86 | Mottled spinefoot | 17 | 1 | 3 | 21 |
| Big roughy | 4 |  |  | 4 | Surgeonfishes | 3 | 1 | 1 | 5 |
| Slimeheads | 3 | 1 | 1 | 5 | Red barracuda | 1028 | 110 | 102 | 1240 |
| Hound needlefish | 4 |  | 1 | 5 | Japanese barracuda | 4469 | 498 | 602 | 5569 |
| Luna lionfish | 2 |  | 1 | 3 | Roudi escolar | 53 | 12 | 8 | 73 |
| Bearded waspfish | 5 |  |  | 5 | Oilfish | 15 | 1 |  | 16 |
| "Mebaru" | 4 | 1 |  | 5 | Gnomefish | 73 | 3 | 9 | 85 |
| Bluefin searobin | 34 | 6 | 6 | 46 | "Kamasu" | 200 | 25 | 35 | 260 |
| Spotted flathead | 4 |  |  | 4 | Largehead hairtail | 8 |  |  | 8 |
| Rogadius | 10 | 1 |  | 11 | Bullet tuna | 3695 | 404 | 470 | 4569 |
| Blackmouth | 72 | 6 | 6 | 84 | Frigate tuna | 857 | 126 | 104 | 1087 |
| Arrow bulleye | 3 |  |  | 3 | "Soudagatsuo" | 402 | 44 | 48 | 494 |
| Half-lined cardinal | 104 | 10 | 11 | 125 | Blue mackerel | 2921 | 268 | 330 | 3519 |
| Japanese bluefish | 3 |  |  | 3 | Chub mackerel | 8118 | 1050 | 1230 | 10398 |
| Common dolphinfish | 29 | 5 | 3 | 37 | Scomber | 755 | 72 | 81 | 908 |
| Giliated threadfish | 2 |  |  | 2 | Pacific bluefin tuna | 2 |  | 1 | 3 |
| Shortfin scad | 12 |  |  | 12 | Shotted halibut (eyed side) | 6 | 1 |  | 7 |
| Japanese scad | 858 | 77 | 89 | 1024 | "Karei" (no-eyed side) | 3 | 1 |  | 4 |
| Amberstripe scad | 62 | 15 | 16 | 93 | Threadsail filefish | 34 | 3 | 3 | 40 |
| Roughear scad | 30 | 3 | 3 | 36 | Half-smooth golden puffer | 132 | 26 | 20 | 178 |
| Decapterus | 59 | 8 | 6 | 73 | "Fugu" | 7 |  | 1 | 8 |
| Whitefin trevally | 85 | 11 | 10 | 106 | Other | 650 | 61 | 147 | 858 |
| Bigeye scad | 461 | 67 | 67 | 595 | Sepia | 5 | 1 | 1 | 7 |
| Greater amberjack | 11 | 5 | 1 | 17 | Swordtip squid | 29 | 6 | 5 | 40 |
| Yellowtail | 525 | 83 | 47 | 655 | Spear squid | 2 |  |  | 2 |
| Japanese jack mackerel | 5378 | 605 | 608 | 6591 | Bigfin reef squid | 40 | 9 | 6 | 55 |
| "Aji" | 22 | 2 | 3 | 27 | Japanese flying squid | 190 | 32 | 27 | 249 |
| Slipmouth | 141 | 17 | 15 | 173 | Luminous flying squid | 8 |  | 2 | 10 |
| "Hiiragi" | 5 |  |  | 5 | Neon flying squid | 4 |  |  | 4 |
| Japanese rubyfish | 2 | 2 |  | 4 | Squid | 15 | 4 | 3 | 22 |
| Rovers | 11 |  |  | 11 | Total | 34602 | 4116 | 4508 | 43226 |

### Training procedure and performance evaluation

The 3909 images were divided into training, validation, and test sets in an 8:1:1 ratio (Table 1). The instance segmentation model adopted with mask-rcnn-R-50-FPN-3x architecture in Detectron2 library (Wu *et al*., 2019) was used for fish detection and classification. This model adopts the Mask R-CNN architecture featuring a ResNet50 backbone and a feature pyramid network (FPN) (Ataullha *et al*., 2023). Furthermore, data augmentation techniques such as random flips (with a probability of 50%) and random resizing, were used to improve model accuracy. The resizing method used was ResizeShortestEdge of the Detectron2 library, which keeps the aspect ratio unchanged, randomly selecting from six values for each input image size (Table 2).

**Table 2.** Model settings for each image size.

| Image size | Input max<br>size train | Input max<br>size test | Input min<br>size test | Input min size train | Anchor generator sizes | Anchor generator<br>strides |
| --- | --- | --- | --- | --- | --- | --- |
| 1333×888 | 1333 | 1333 | 800 | (640, 672, 704, 736, 768, 800) | (32, 64, 128, 256, 512) | (4, 8, 16, 32, 64) |
| 2000×1333 | 2000 | 2000 | 1200 | (960, 1008, 1116, 1104, 1152, 1200) | (48, 96, 192, 384, 768) | (6, 12, 24, 48, 96) |
| 2666×1777 | 2666 | 2666 | 1600 | (1280, 1344, 1408, 1472, 1536, 1600) | (64, 128, 256, 512, 1024) | (8, 16, 32, 64, 128) |

The input sizes of the images were set to the default size (1333×888), 1.5 times both sides (2000×1333), and twice the size (2666×1777), as the default image size for the Mask R-CNN model in the Detectron2 library is 1333 pixels on the long side. The model settings for image size are shown in Table 2. Each dataset was used for training using the weights of the pretrained model (fine-tuning). The number of iterations was 270,000, and the learning rate was set to 0.01. The batch size was set to four. The model with 270, 000 iterations was used for this study since the total loss did not increase and was consistently low (Supplemental Table 2).

The development environment was Python3.9.13 and Pytorch1.10.0+cu102. The GPU was NVIDIA Quadro RTX 8000.

### Performance evaluation

To evaluate model performance, the mean average precision (mAP), which is often used for object detection (Mao *et al*., 2023), was adopted. Precision and recall were calculated as follows:

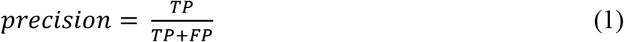

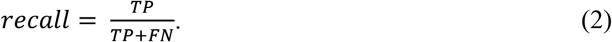

If the predicted class matched the true class, the prediction was marked as true positive (TP). If the class did not match, the ground truth was marked as false negative (FN) and the corresponding network prediction as false positive (FP). All network predictions that did not have an associated ground-truth mask were marked as FPs, and all ground truth masks that did not have an associated network prediction were marked as FN.

A trained model generates a TP using the coordinates of the mask and confidence score (the confidence for each detection mode by model) (Redmon *et al*., 2016). Average precision (AP) is the area under the precision-recall curve and is calculated as follows (Butt *et al*., 2023):

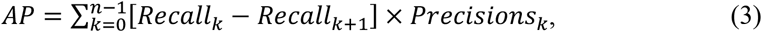

where *k* is the number of the objects ordered by confidence score.

To calculate mAP, the mean value of all the AP classes is taken, as follows:

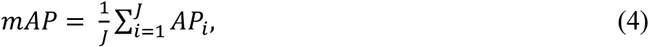

where *J* is the total number of classes with *i* representing each class.

To determine the precision-recall curve, the true and false positive values of the prediction must be calculated. To determine the success of detection, we used intersection of union (IoU), defined as follows (Tang *et al*., 2020):

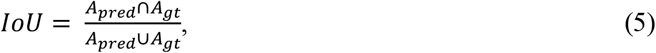

where A_pred_ and A_gt_ represent the areas in the predicted and ground truth masks, respectively. Then, we set a threshold for IoU, for example 0.5, such that if the IoU exceeded the threshold, the detection was marked as correct. Multiple detections of the same object were considered as a single correct detection and the others as false detections. After obtaining the true-positive and false-positive values, the precision-recall curve and mAP were calculated (Tang *et al*., 2020). The mAP was calculated with IoU being incremented from 0.50 to 0.95 in increments of 0.05, and the average (mAP50-95) was defined as

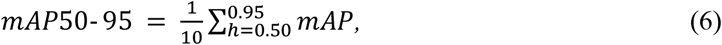

for *h* in the range of 0.50 to 0.95 in increments of 0.05 (total number of 10). This index was used to compare the detection accuracies of the models trained for each image size.

To examine the required number of fish per class for high-accuracy classification, the relationship between the number of fish per class and AP was plotted using scatter plots.

### Evaluation of factors affecting the improvement of species identification accuracy with changes in image size

A generalized linear mixed model (GLMM) was used to analyze the effect on accuracy. First, three AP50_*i,j*_ (AP values for IoU of 0.50 and *j* = 1,2,3) were calculated for each class *i*. Here, AP50_*i,j*=1_ represents the score obtained using the smallest input image size for class *i* (1333×888); AP50_*i,j* =2_ represents the score obtained using the second smallest input image size (2000×1333); and AP50_*i,j* =3_ represents the score obtained using the largest input image size (2666×1777). Next, the differences *D*_*i,j*_ were calculated based on AP50_*i,j*=1_ for each class *i* as follows:

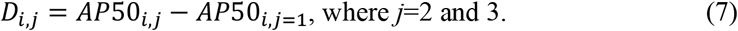

These differences indicate the extent of performance improvement, as measured by AP50, for increases in image size although only two values were calculated (i.e., *D*_*i,j*=2_ and *D*_*i,j*=3_) for each class *i*. The *D*_*i,j*_ were assumed to follow the following model:

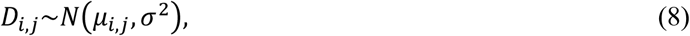

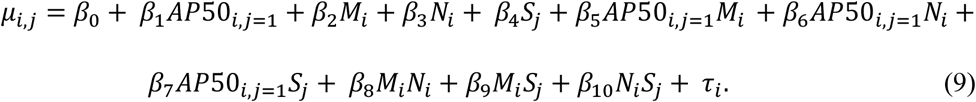

where *μ* and *σ*^2^ are the mean and variance of a Gaussian distribution, respectively; *β*_0_ is the intercept; and *β*_1_ ∼*β*_10_ are the coefficients of explanatory variables. M is the average area (pixels) in each class with the original image size as (3200×4800); N is the number of training data samples; and S is the total number of pixels for each image size, calculated as the difference from the smallest image size. The second interaction terms between each combination were also used as explanatory variables. The term *τ*_*i*_ represents the random effect of each class *i*. The analysis was conducted using R version 4.2.2 (R Core Team, 2022) with Package lme4. The explanatory variables were selected using exhaustive research based on Akaike’s information criterion (AIC), and the model with the lowest AIC was chosen as the best. Classes and image sizes for which the AP values could not be calculated because of a lack of test data or no prediction, were excluded from the analysis.

## Results

In the experiments, only 62 of the 85 classes could be evaluated for accuracy due to the heterogeneity in the number of training and test datasets. This heterogeneity occurred because the photos per individual fish were not divided into training and test data, but rather per photo. The mAP50-95, which is a measure of object detection and classification accuracy, was 0.580 or better for all image sizes (Table 3). The mAP tended to improve with a relatively large image size, obtaining the best mAP50-95 for a size of 2000×1333 (Table 3). The maximum improvement was 0.025, which was the difference between an mAP50-95 of 0.605 at 2000×1333 and 0.580 at 1333×888 (Table 3). AP50 was 0.95 or better for Japanese jack mackerel (*Trachurus japonicus*), yellowtail (*Seriola quinqueradiata*), and crimson sea-bream (*Evynnis tumifrons*) at an image size of 2666×1777, except for some classes with AP50 of 1.00 probably due to insufficient test data (Table 4). A trend toward increased accuracy was not observed as the amount of data increased, but the variance in accuracy tended to decrease with an increase in the data (Figure 2). However, blue mackerel had a low accuracy of approximately 0.5, even with more than 3,000 data. Classes containing multiple species, such as “Aji,” “Iwashi,” Decapterus, “Kamasu”, scomber, and squid, tended to have low accuracy.

**Table 3.** Accuracies (mAP) by IoU for each image size.

| Image size | IoU |  |  |  |  |  |  |  |  |  |  |
| --- | --- | --- | --- | --- | --- | --- | --- | --- | --- | --- | --- |
|  | 0.5 | 0.55 | 0.6 | 0.65 | 0.7 | 0.75 | 0.8 | 0.85 | 0.9 | 0.95 | 0.50:0.95 |
| 1333×888 | 0.733 | 0.729 | 0.714 | 0.693 | 0.683 | 0.655 | 0.617 | 0.544 | 0.386 | 0.041 | 0.580 |
| 2000×1333 | 0.747 | 0.745 | 0.743 | 0.724 | 0.715 | 0.677 | 0.642 | 0.590 | 0.415 | 0.047 | 0.605 |
| 2666×1777 | 0.741 | 0.738 | 0.721 | 0.706 | 0.683 | 0.668 | 0.650 | 0.577 | 0.390 | 0.057 | 0.593 |

**Table 4.**
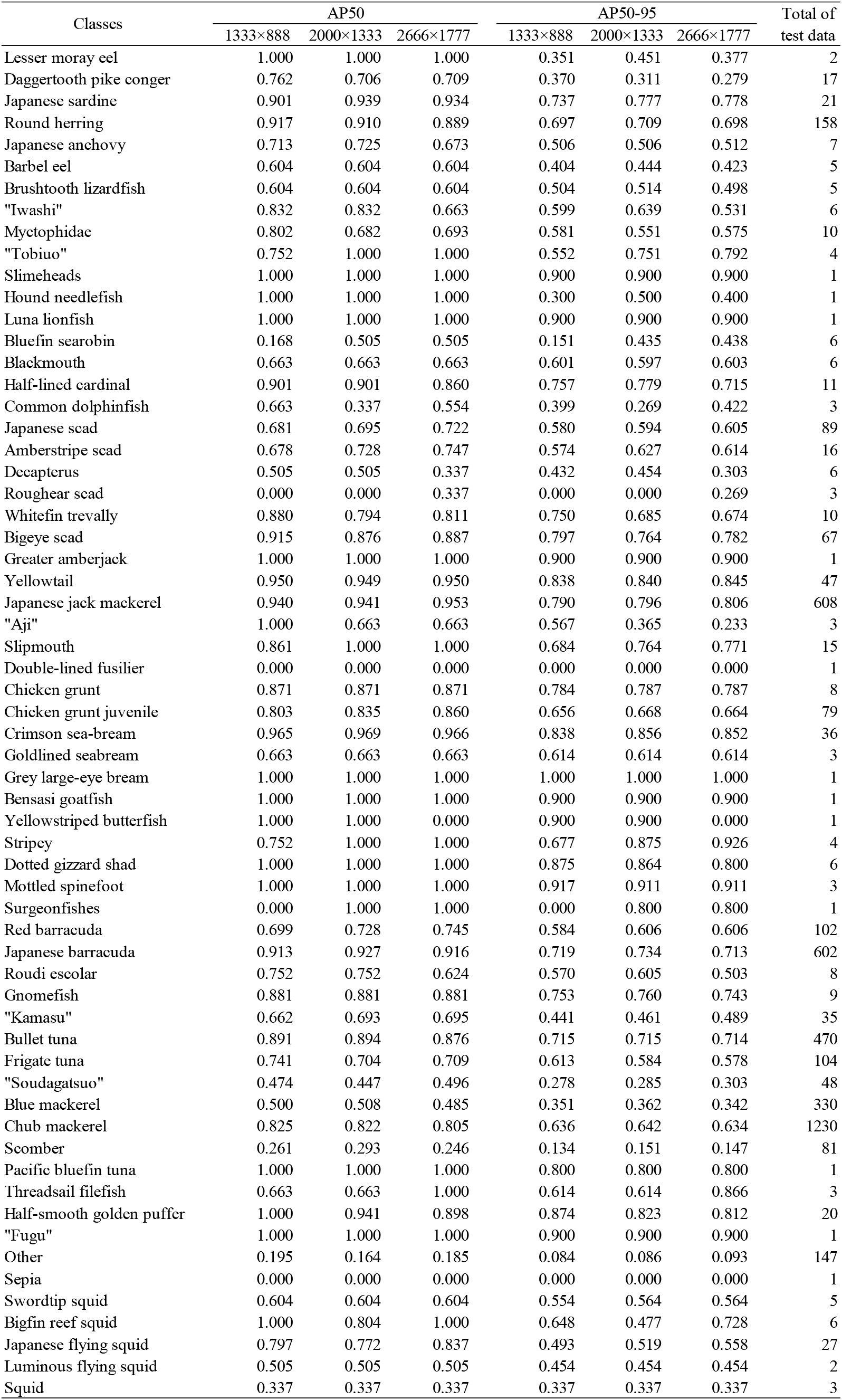
AP50 and AP50-95 by classes for each image size.

**Fig. 2.**
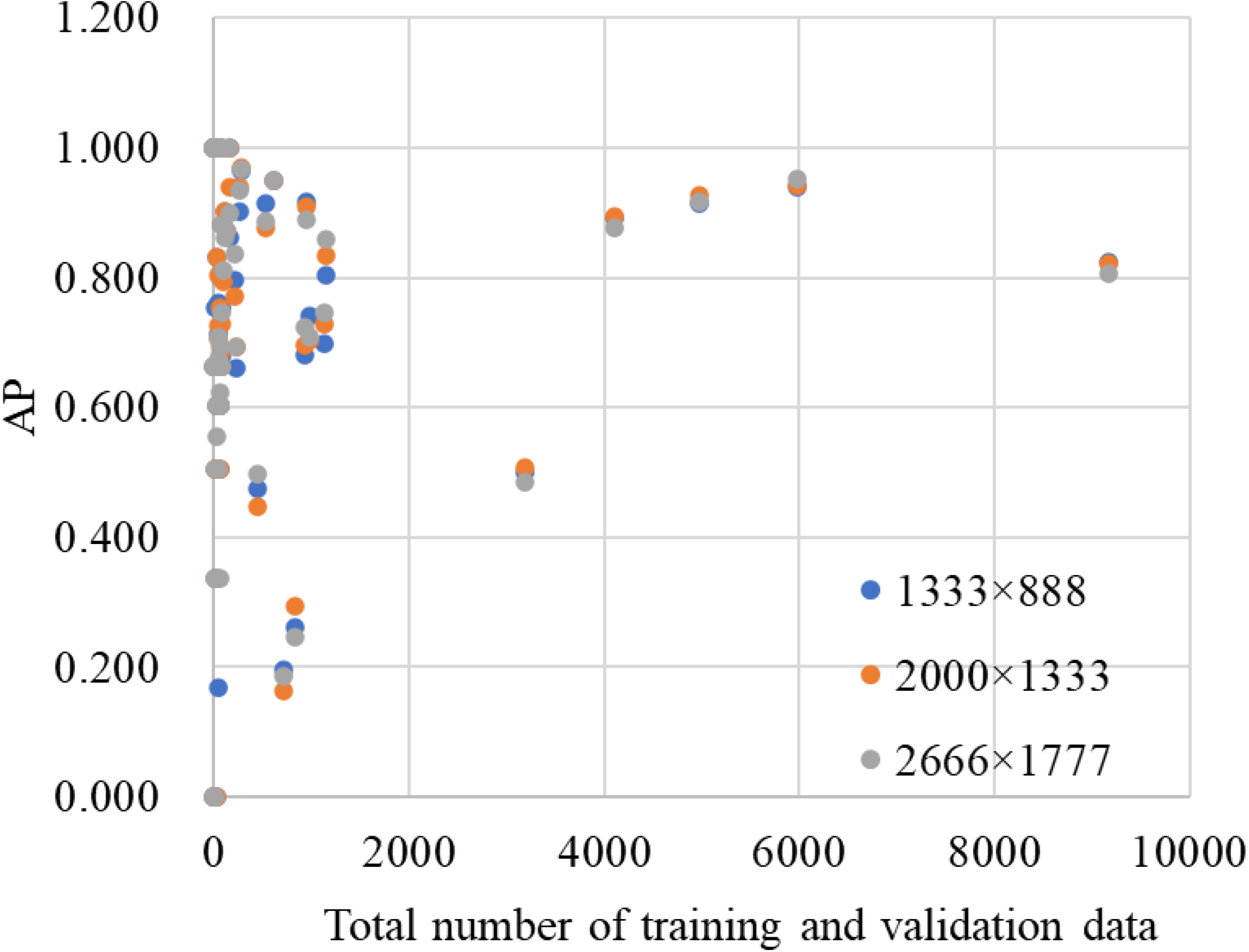
Scatter plots of the relationship between the number of data and AP.

As a result of modeling the AP50 difference using GLMM (Table 5), the AP50 at an image size of 1333×888 was the only explanatory variable in the best model, and accuracy tended to improve with lower AP50 values for the 1333×888 image size (Figure 3). Average area by class was not selected as an explanatory variable. However, the accuracy on blue mackerel did not improve despite displaying a low AP50 for the 1333×888 image size (Table 4).

**Table 5.** Details of the best model. Only fixed effects were indicated. *P value < 0.05.

| Explanatory variables | Estimate | Standard error | df | t value | p value |
| --- | --- | --- | --- | --- | --- |
| Intercept | 0.204 | 0.055 | 60.000 | 3.721 | 4.40E-04 * |
| AP50<sub>*ij=1*</sub> | -0.263 | 0.070 | 60.000 | -3.766 | 3.80E-04 * |

**Fig. 3.**
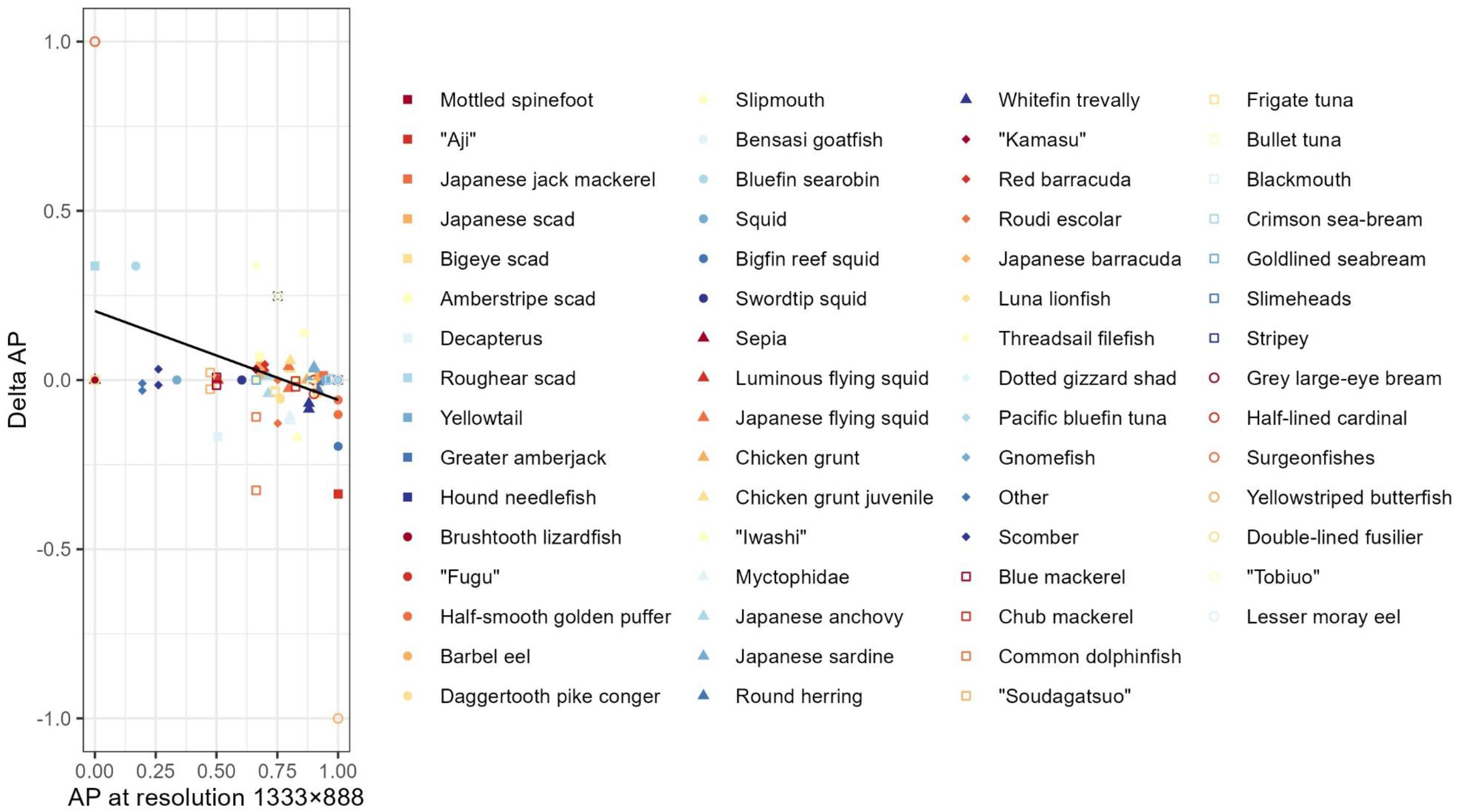
The relationship between AP at minimum image size and the difference between AP at minimum image size and AP at larger image size. Points indicate measured values and lines indicate predicted lines modeled by GLMM.

## Discussion

### Effect of changing image size

In general, higher image size increased classification accuracy (Huang *et al*., 2018), resulting in improvements in some of the previous studies (Camgözlü and Kutlu, 2020; Gope and Fukai, 2022). In our study, the detection and classification accuracy also tended to improve when the image size was larger. However, the most accurate results were not necessarily obtained with the model trained using the largest image size; the model trained on 2000×1333 images was more accurate.

A possible reason for the model trained on the largest image size not being the most accurate, could be the effect of the number of pretraining weights. The model trained without pretrained weights, which was examined a priori, recorded the best mAP50-95 at the largest size of 2666×1777, but the overall mAP was lower than that of the fine-tuning model (Supplemental Table 2). Therefore, the finding that the fine-tuning model trained on the largest image size was not the most accurate may be due to the effect of the pretrained weights, which were trained on the default image size. Tan and Le (2019) reported that changing the number of layers along with the depth of the network and image size, rather than just the image size, is useful in improving classification accuracy. This may also explain why the largest image size did not result in the highest mAP50-95 in our study, although mAP is an evaluation index of object detection and classification.

Classes with low classification accuracy for the smallest image size tended to improve the classification accuracy with increasing image size (Table 5). However, the accuracy tended not to change with the average area of each class. There may have been no classes small enough to affect the accuracy of detection and classification.

In our study, the accuracy improvement was 0.014 for mAP50-95 and 0.008 for mAP50 compared to those of maximum image size and default size. On the other hand, the computation time was 3.4 times longer for inference and 2.7 times longer for training (Supplemental Table 3). The image size used in practice is a trade-off between the required accuracy and computational speed for real-time data. A stock assessment is usually conducted once a year; therefore, long computation times may not be a significant issue.

### Application to stock assessment

In this study, object detection and classification were performed on 85 classes using deep learning approaches, and their accuracy was evaluated on 62 classes. Although previous studies using deep learning approaches have already identified fish species by taking images of the catches onboard using several fishing methods (Lu *et al*., 2020; Lekunberri *et al*., 2022), our study is the first to identify fish species caught by set-nets, which have more fish species than most of the other methods (Ministry of Agriculture Forestry and Fisheries, 2023). The number of classes in our study was approximately twice as large as the maximum number of 31 species reported previously (Mei *et al*., 2021). By fixing the camera on the conveyor belts at fishing ports, length information can be obtained from fishing methods that were previously not reported for acquiring image data on board. Our study expands the number of fishing methods and target species for which information on fish length can be obtained through deep learning.

Some fish species, such as the Japanese jack mackerel, yellowtail, and crimson sea-bream, were predicted particularly accurately, with AP values of almost 0.95 or higher. The prediction for Japanese jack mackerel was highly accurate even in the presence of similar species classes and classes with multiple species. The high accuracy of yellowtail and crimson sea-bream might be because of the small number of very similar species such as yellowtail kingfish (*Seriola aureovittata*) and red sea-bream (*Pagrus major*). If similar fish species are present in the catches, the classification accuracy is more likely to decrease. However, these species are ready to be used for length information in stock assessments in the ports where similar fish species are not caught although the accuracy of the classification data used for stock assessment is determined through stakeholder agreement.

### Effect of number of data samples and quality of data on accuracy

Deep learning requires a massive amount of data ranging from 5,000 to 10,000 samples per label(Goodfellow *et al*., 2016). In our study, there was no linear positive relationship between the number of data and classification accuracy. Among the species with a small amount of data, some were identified with high accuracy and others with low accuracy. However, for those with an AP50 of 0.80 or higher, the number of training and test data was more than 4,000. The accuracy of identifying blue mackerel was extremely poor at 0.5, even with approximately 3,000 data. One possible reason for poor accuracy is mislabeled training data. There are two identification characteristics that distinguish between chub and blue mackerels: (i) the spotted color pattern on the underbody and (ii) the ratio of the fork length to the length of the basal portion of the first dorsal fin on the first to ninth spine rays (Nakabo, 2013). In this study, only the spotted color pattern was used to identify the mackerels and label the training data, since the second characteristic could not be confirmed on images. However, the first characteristic varies greatly among individual fish, making it extremely difficult to distinguish chub mackerel from blue mackerel on the images based only on the first characteristic. In addition, manually adjusted lights may complicate identification. Depending on the position of the fish in the images, the light sometimes strongly reflects off from the body of the fish. These reflections may obscure the spotted pattern of the blue mackerel, resulting in significant degradation in the quality of the annotation labels. To solve this problem, the lighting probably needed to be optimized by installing special boxes for imaging (van Essen *et al*., 2021) or by bouncing of the light (Hunter *et al*., 2021). Additionally, the spotted pattern has a gradation, which makes identification difficult using only the first character difficult. They can be classified more correctly using the second characteristic, but it is difficult to check the character on the images while annotating and labeling at the desk. To examine whether these species can be identified using deep learning approaches, images of fish that were correctly identified by the second identification characteristic using real fish, should be collected.

Although the accuracy of classes including multiple species tended to be higher for group-level classifications (Supplemental Tables 1 and 4), classes that included multiple species could not be identified at the species level, tending to display low accuracy for species-level classifications. This study attempted to classify fish at the species level for stock assessment. However, identifying all fish species only from images taken on a conveyor is difficult; therefore, we created classes that included multiple species. When species-level identification is conducted using images of fish catches, some fish are probably not identifiable at the species level, as in the present study. The low accuracy of these classes is not a serious problem because multiple species classes are not used in actual stock assessments; however, the impact of these classes on the accuracy of other classes needs to be examined. Because the number of multiple-species class data in this study was smaller than that of single-species class data, the accuracy of single-species classes was less affected by multiple-species class fish being misclassified as other single-species classes. This impact may be greater with a larger number of multiple-species class data. The problem of being unable to identify species may always occur in the future when creating training data at the species level from photographs of catches, which will necessitate some improvement in the future.

## Conclusion

This study classified 85 classes of fish using images of set-net catches that landed on a conveyor belt in a fishing port. The models were trained on different image sizes to evaluate the change in accuracy of detection and classification of fish species, thereby observing the trend that the larger the image size, the better the accuracy. However, the accuracy was not the highest for the largest image size, suggesting that fine-tuning decreases accuracy when image size is significantly different from the default image size. A trend of improved accuracy was observed for species that had a low accuracy at the smallest image size, and the proposed method was considered useful as a simple way of improving accuracy. Deep learning approaches have never been applied to images taken on a conveyor belt in a fishing port, and our study can help obtain fish length information on a larger number of fish species for precise stock assessment.

## Aknowledgement

We acknowledge Mr. Tomoyuki Kuwahara, Dr. Toru Kitamura, Ms. Satoko Tamura, Mr. Ken Nakagawa for their assistance in obtaining fish images. Dr. Yutaka Osada worked with us to acquire images of the fish and provided helpful comments early in our study. Moreover, the authors would like to thank Editage (www.editage.com) for English language editing and TTPM Inc. for providing high-quality annotated data.

## Author contributions

Conceptualization: YI, and YS; Data Curation: YI, YS, and MM; Formal Analysis: YI; Funding: YS; Investigation: YI, and YS; Methodology: YI, YS, NT, RK, and HY; Supervision: YS; Writing— original draft: YI; Writing—review and editing: YI, YS, MM, TN, RK, and HY.

## Data availability

The authors do not have permission to share data.

