## Supplementary material captions for "Effects of input image size on the accuracy of fish identification using deep learning"

Supplemental Table 1. List of fish classes their scientific names, and groups.

Supplemental Table 2. Detection accuracy (mAP) for each image size using non-fine tuning model. Only the learning rate setting was changed from 0.01 to 0.001 as training was not possible.

Supplemental Table 3. Computation time for training and inference.

Supplemental Table 4. AP50 and AP50-95 by class. Each average value is the same as that of mAP50 and mAP50-95.

Supplemental Figure 1. Confusion matrices for each image of size: (a) 1333×888, (b) 2000×1333, (c) 2666×1777.

Supplemental Figure 2. Loss for each image of size: Fine-tuning models, (a1) 1333×888, (a2) 2000×1333, (a3) 2666×1777. Non-fine tuning models, (b1) 1333×888, (b2) 2000×1333, (b3) 2666×1777.

Supplemental Figure 3. Example of Japanese multiple species classes. Classes not evaluated due to lack of test data were excluded. (a-1) "Iwashi“ class. Either of the following species; *Sardinella aurita*, *Sardinops melanostictus*, *Etrumeus micropus*, *Engraulis japonica*, Myctophidae. (a-1) "Iwashi“ class. Either of the following species; *Sardinella aurita*, *Sardinops melanostictus*, *Etrumeus micropus*, *Engraulis japonica*, Myctophidae. (b) "Tobiuo“ class. Either of the following species; Exocoetidae. (c) "Aji “ class. Either of the following species; *Alectis ciliaris*, *Atropus hedlandensis*, *Caranx papuensis*, *Caranx sexfasciatus*, *Decapterus akaadsi*, *Decapterus macarellus*, *Decapterus macrosoma*, *Decapterus maruadsi*, *Decapterus muroadsi*, *Decapterus tabl*, *Decapterus* spp., *Selar crumenophthalmus*, *Trachurus japonicus*, *Uraspis helvola*. (d) “Kamasu“ class. Either of the following species; *Sphyraena pinguis*, *Sphyraena* sp., *Promethichthys Prometheus*. (e-1) “Soudagatsuo“ class. Either of the following species; *Auxis rochei rochei*, *Auxis thazard thazard*. (e-2) “Soudagatsuo“ class. Either of the following species; *Auxis rochei rochei*, *Auxis thazard thazard*. (f) “Fugu “ class. Either of the following species; Tetraodontidae.
