## Supplementary figures and images for "Effects of input image size on the accuracy of fish identification using deep learning"

### Supplemental_figure_1a_confusion_matrix_new_name_1333.png

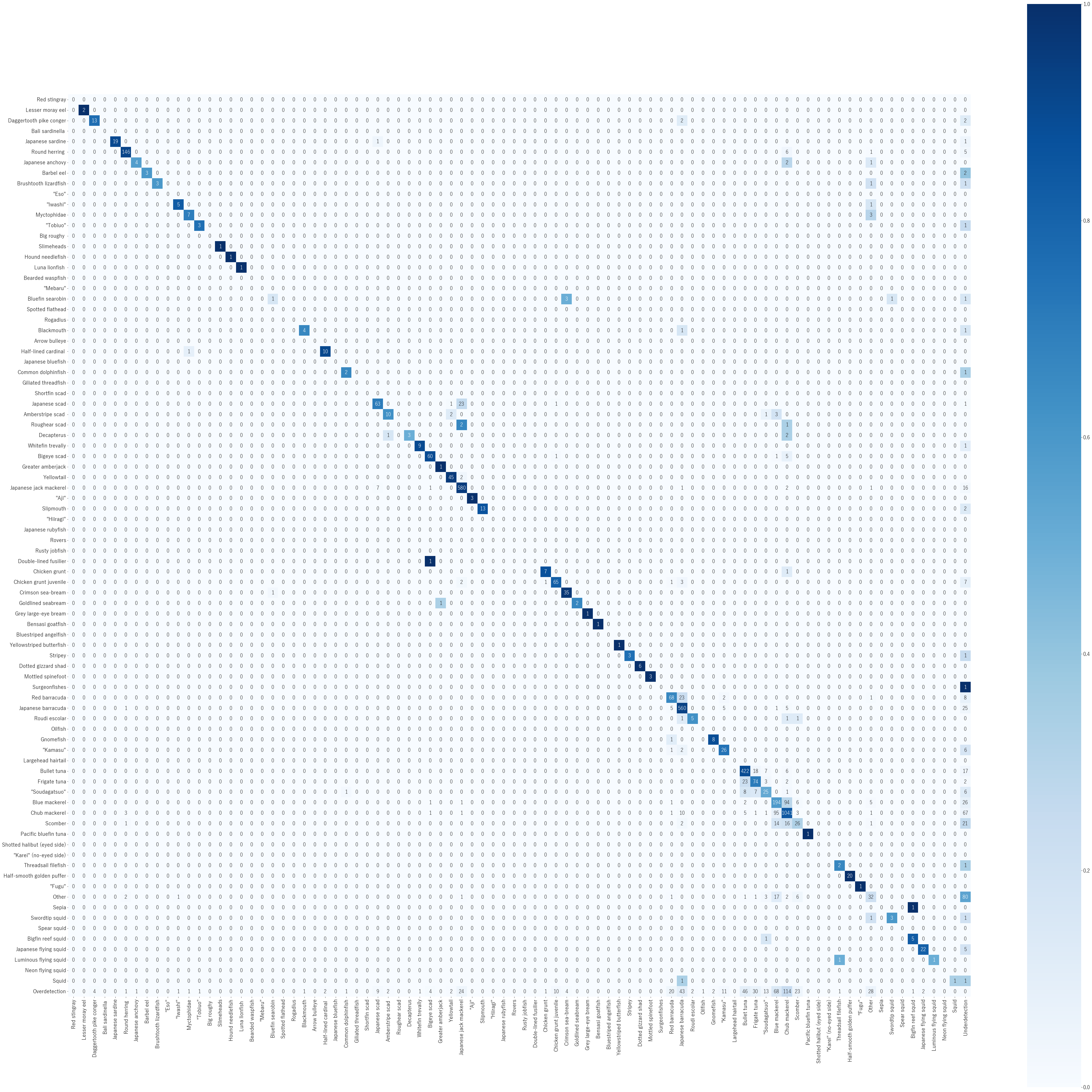

### Supplemental_figure_1b_confusion_matrix_new_name_2000.png

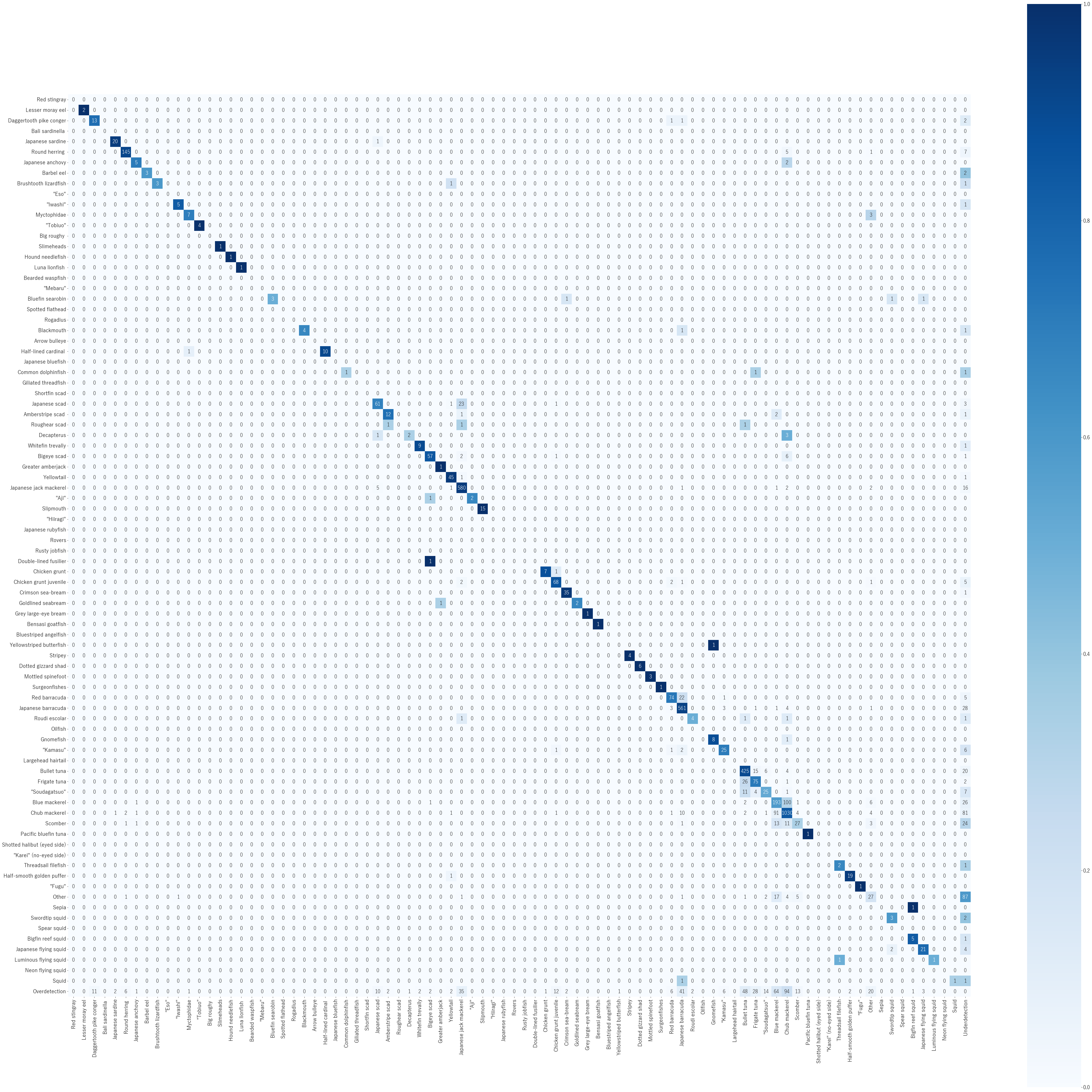

### Supplemental_figure_1c_confusion_matrix_new_name_2666.png

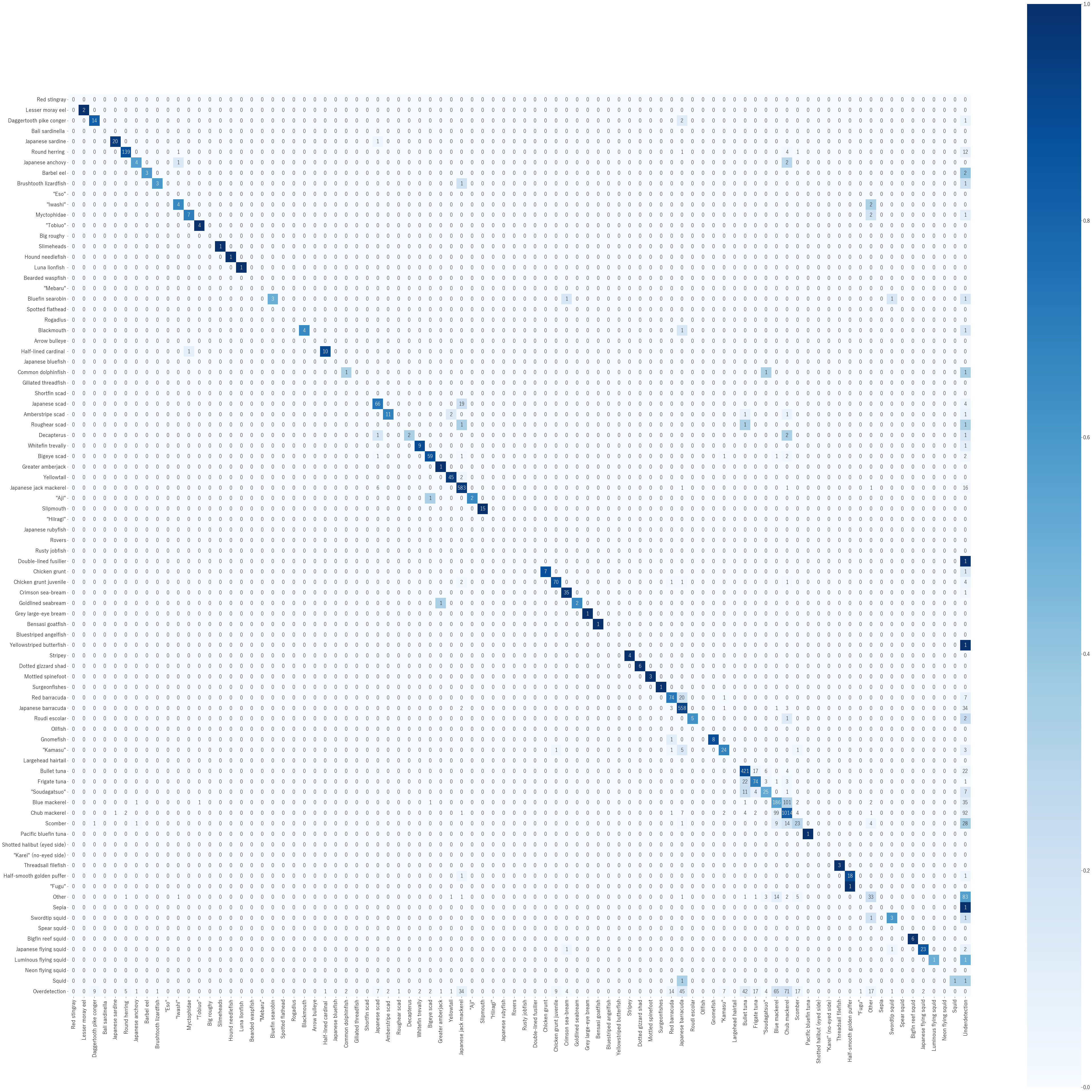

### Supplemental_figure_2_a1_loss_1333.jpg

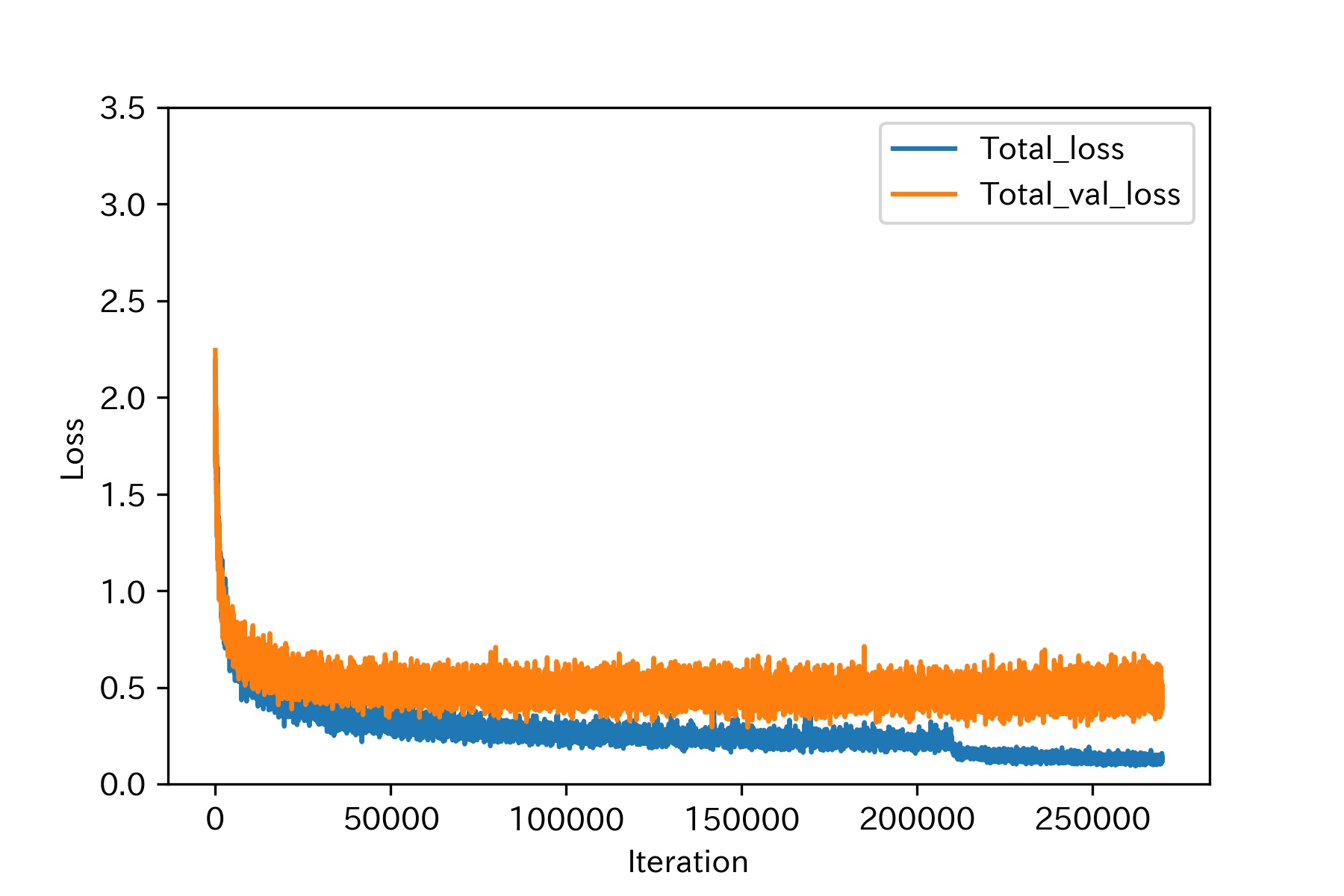

### Supplemental_figure_2_a2_loss_2000.jpg

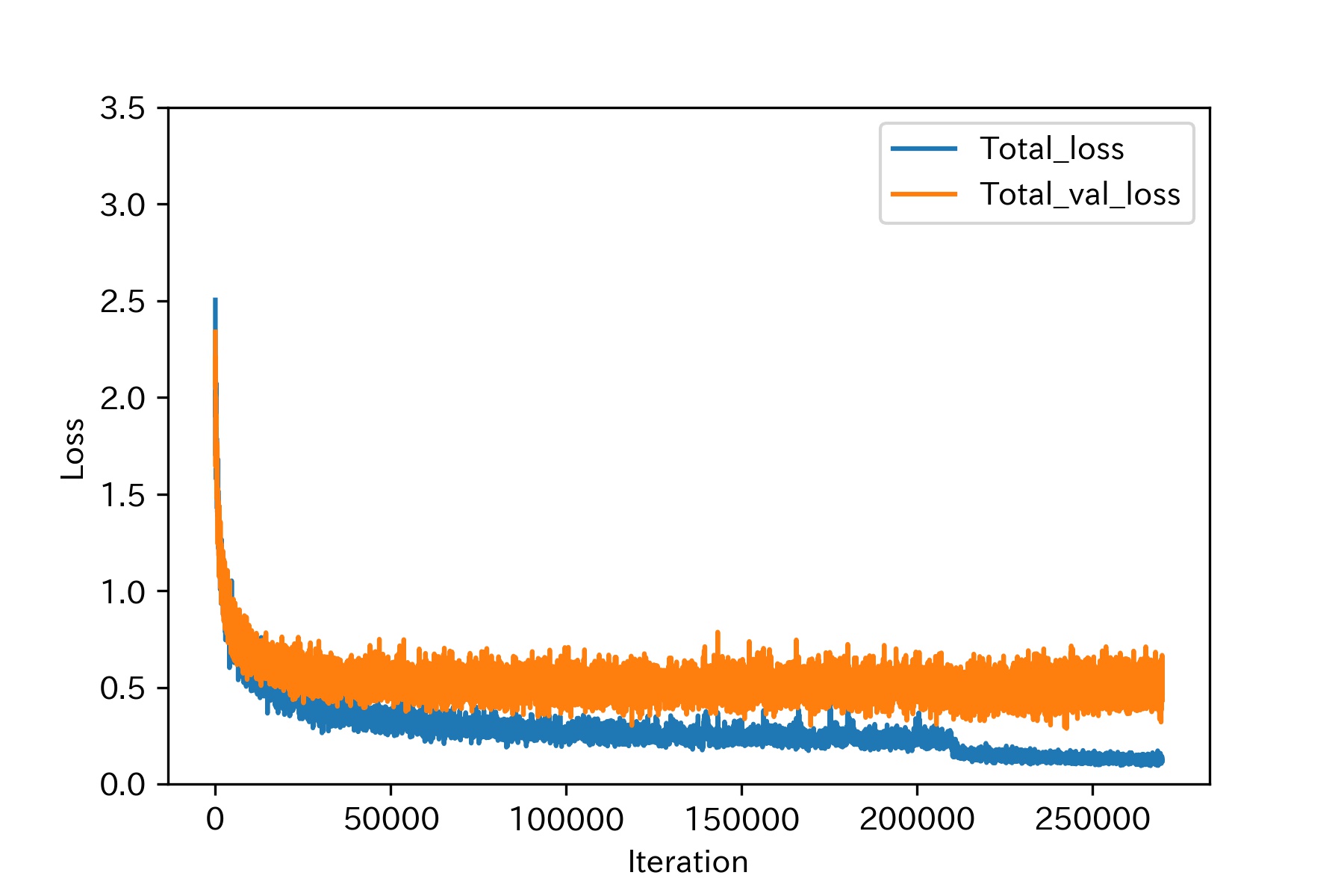

### Supplemental_figure_2_a3_loss_2666.jpg

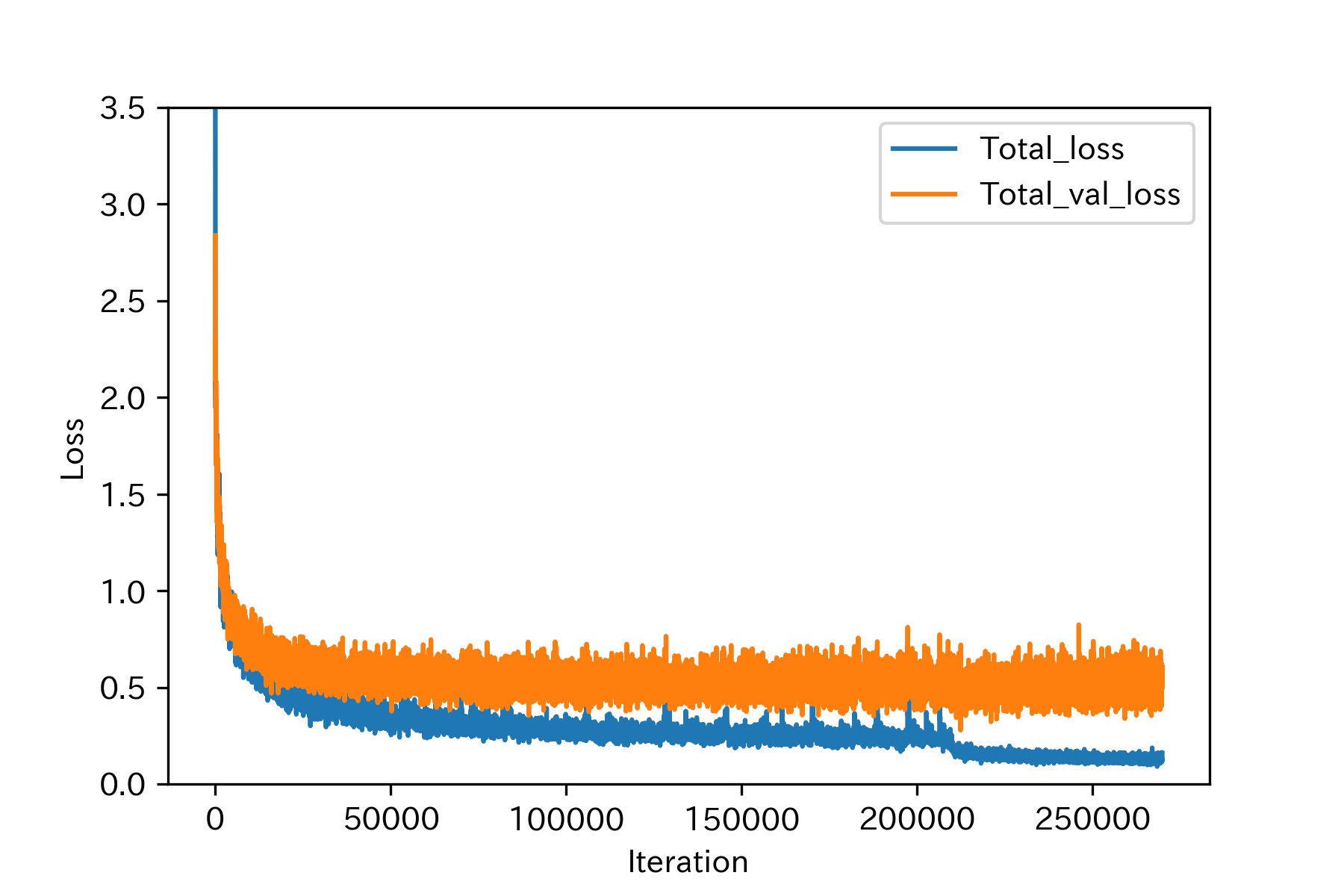

### Supplemental_figure_2_b1_loss_1333_now.jpg

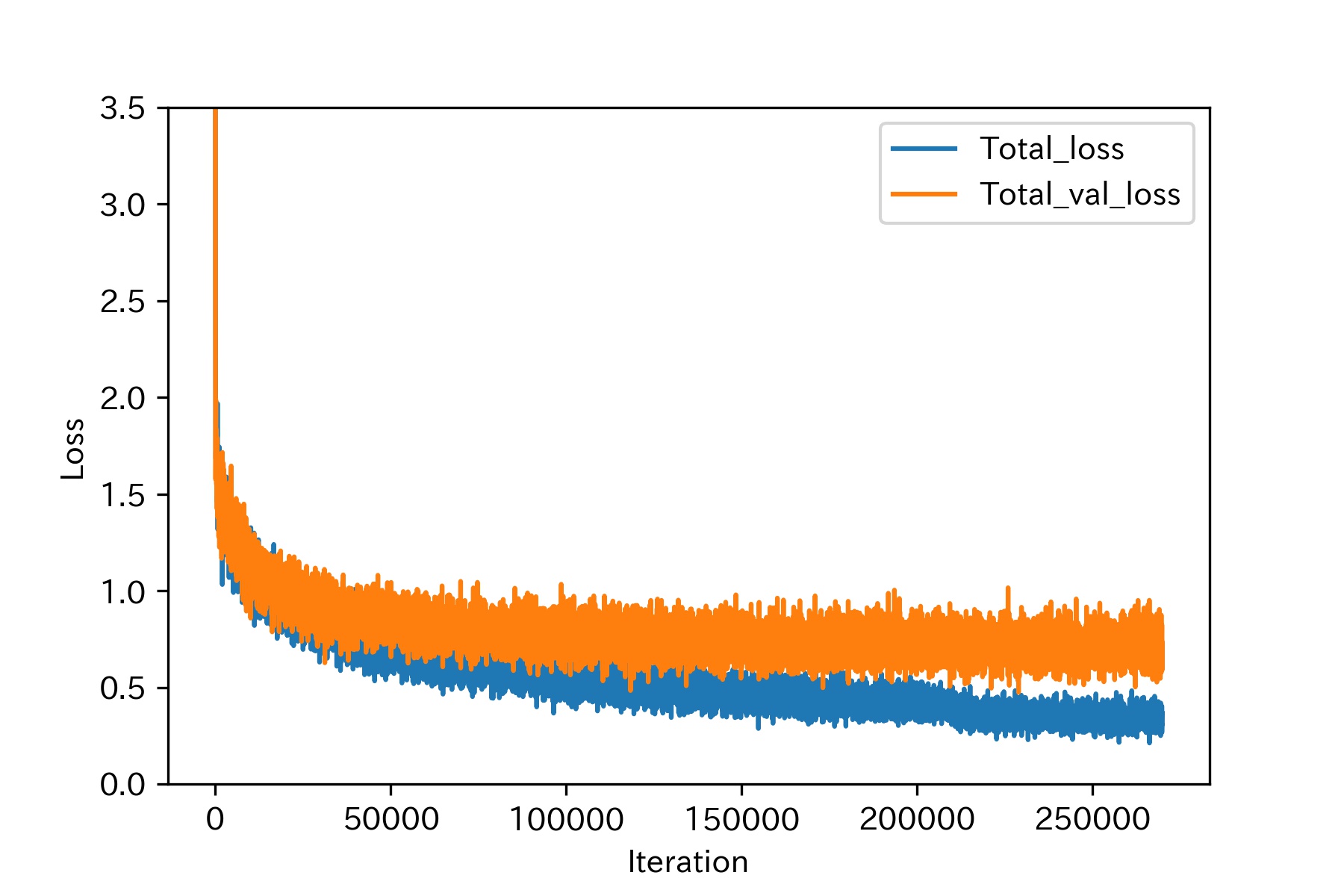

### Supplemental_figure_2_b2_loss_2000_now.jpg

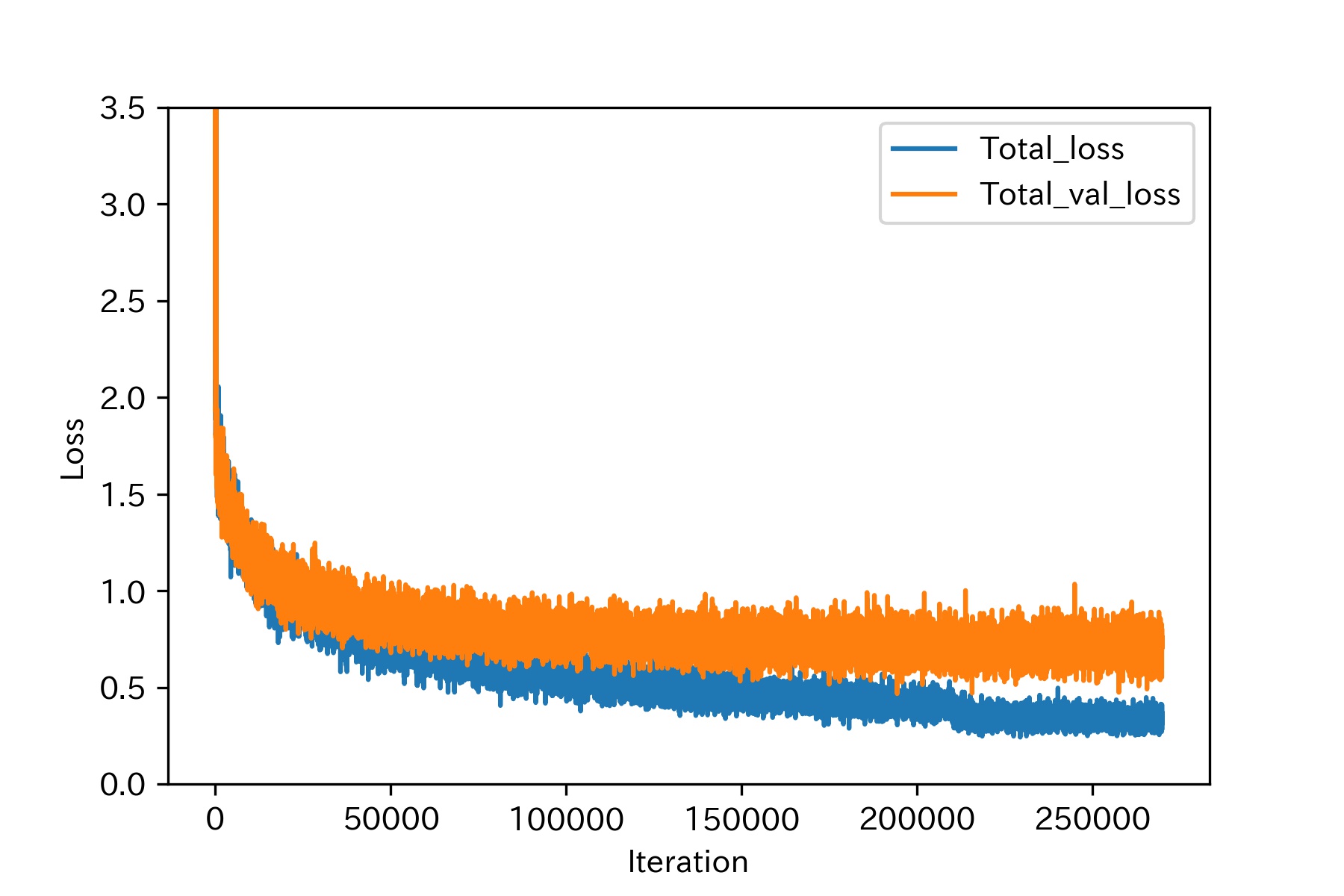

### Supplemental_figure_2_b3_loss_2666_now.jpg

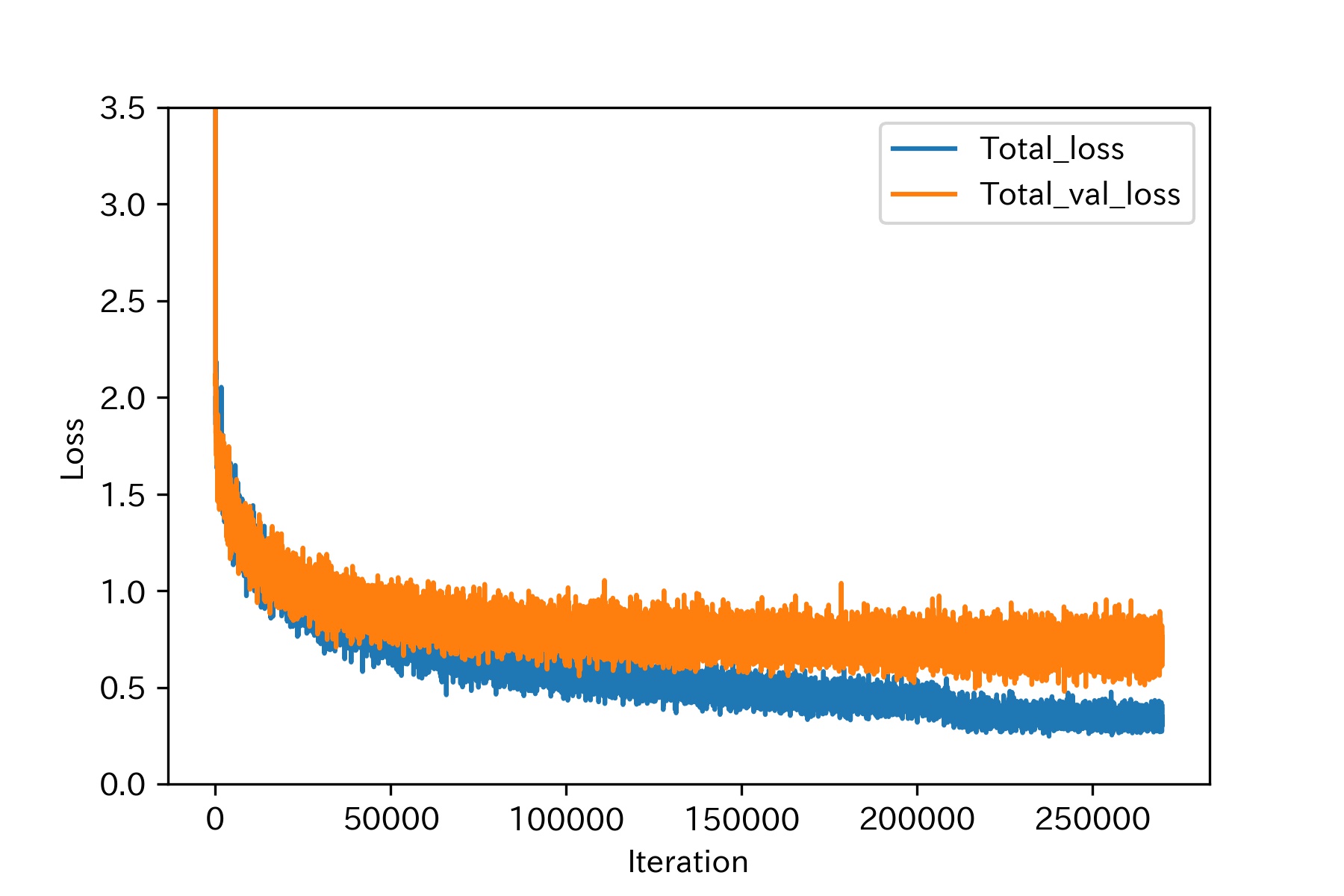
