## Supplementary figure 3 for "Effects of input image size on the accuracy of fish identification using deep learning"

(a-1)

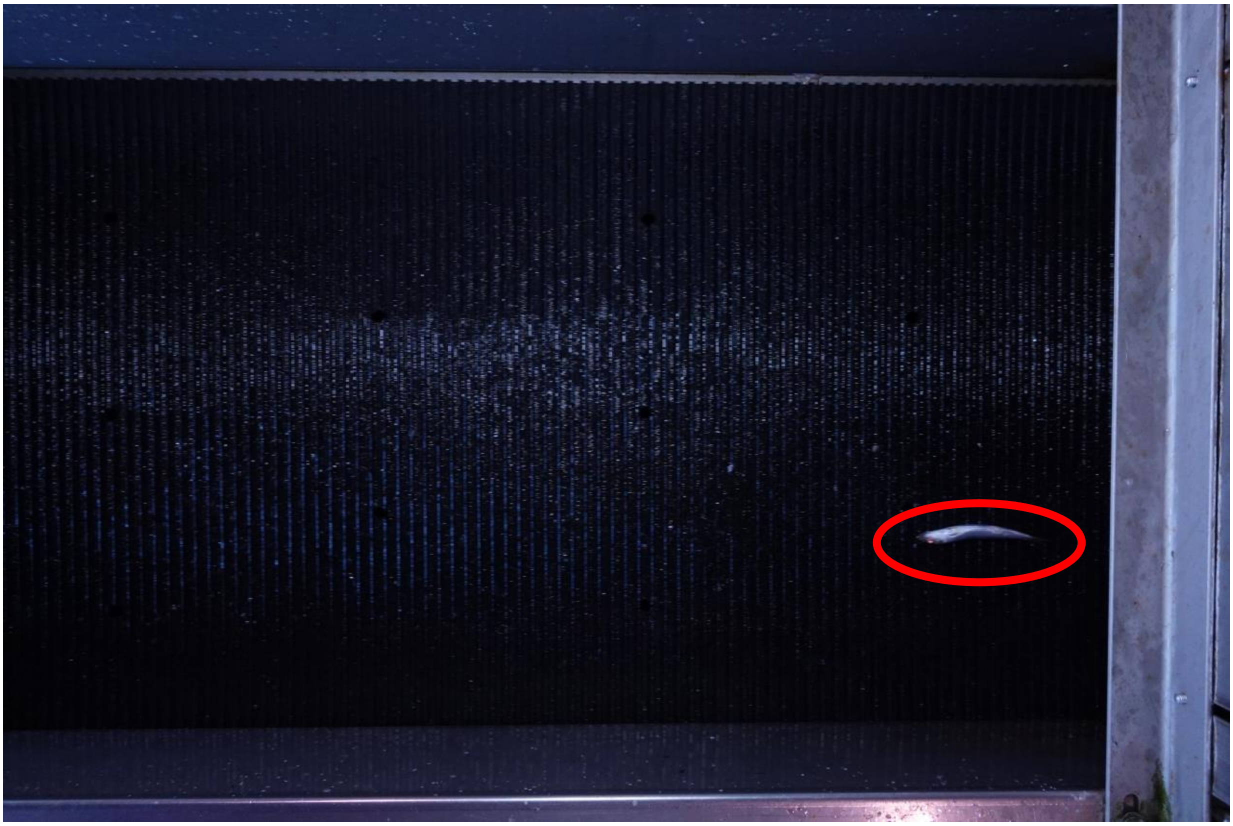

(a-2)

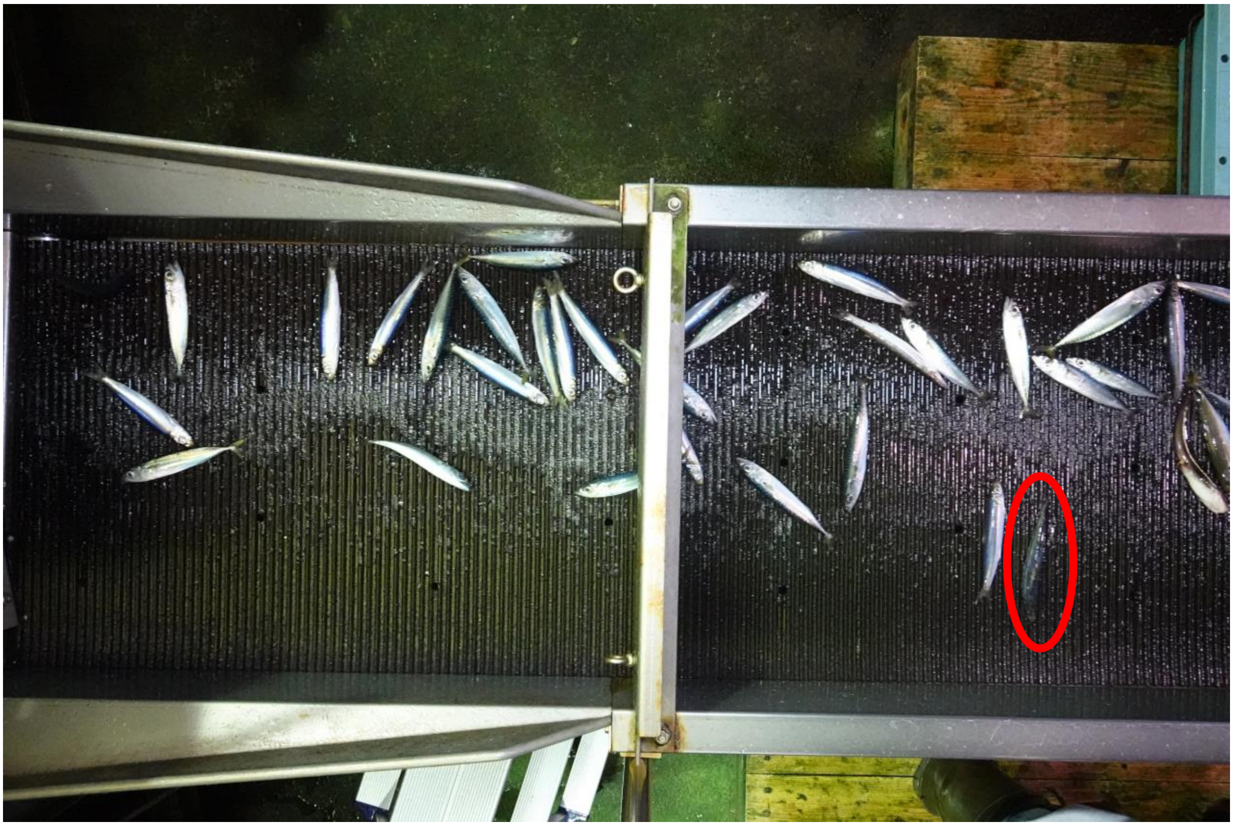

Supplemental Figure 3. Example of Japanese multiple species classes. Classes not evaluated due to lack of test data were excluded. (a-1) "Iwashi“ class. Either of the following species; *Sardinella aurita*, *Sardinops melanostictus*, *Etrumeus micropus*, *Engraulis japonica*, Myctophidae. (a-1) "Iwashi“ class. Either of the following species; *Sardinella aurita*, *Sardinops melanostictus*, *Etrumeus micropus*, *Engraulis japonica*, Myctophidae.

(b)

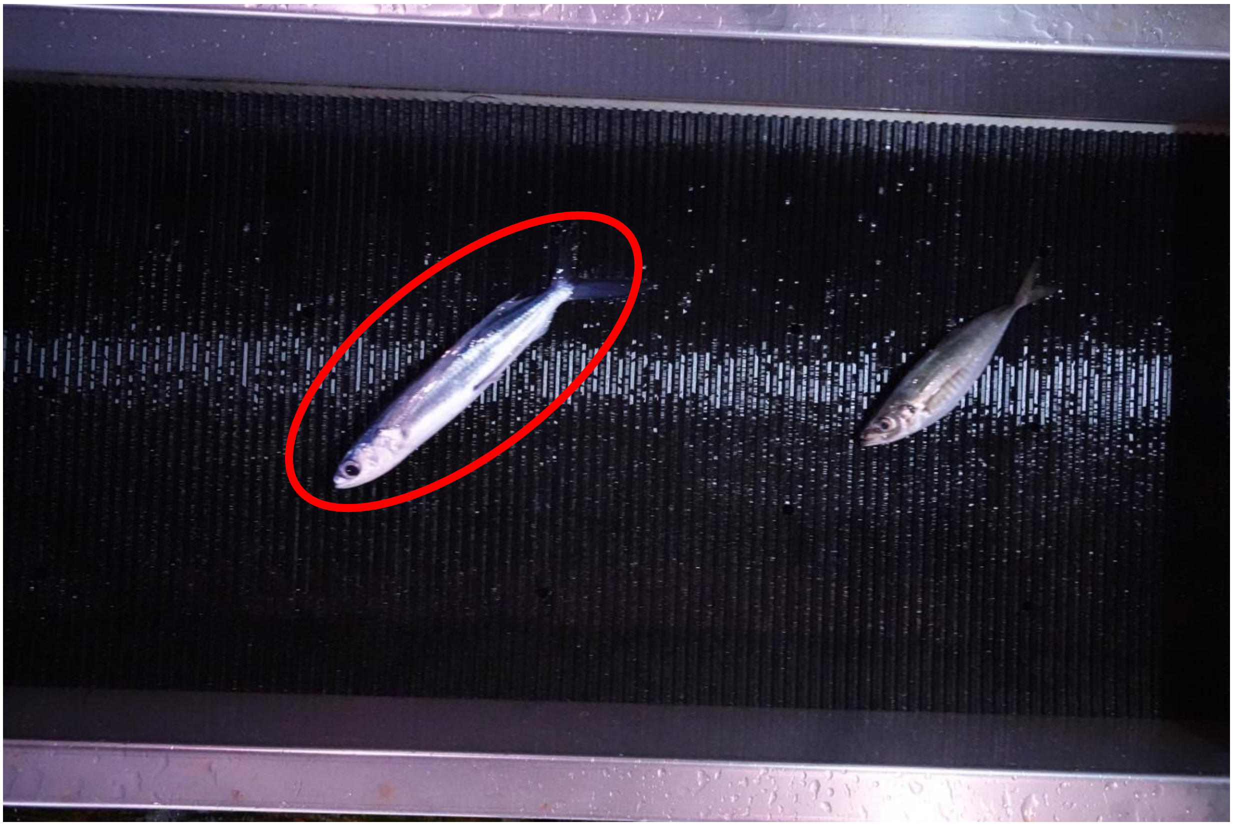

(c)

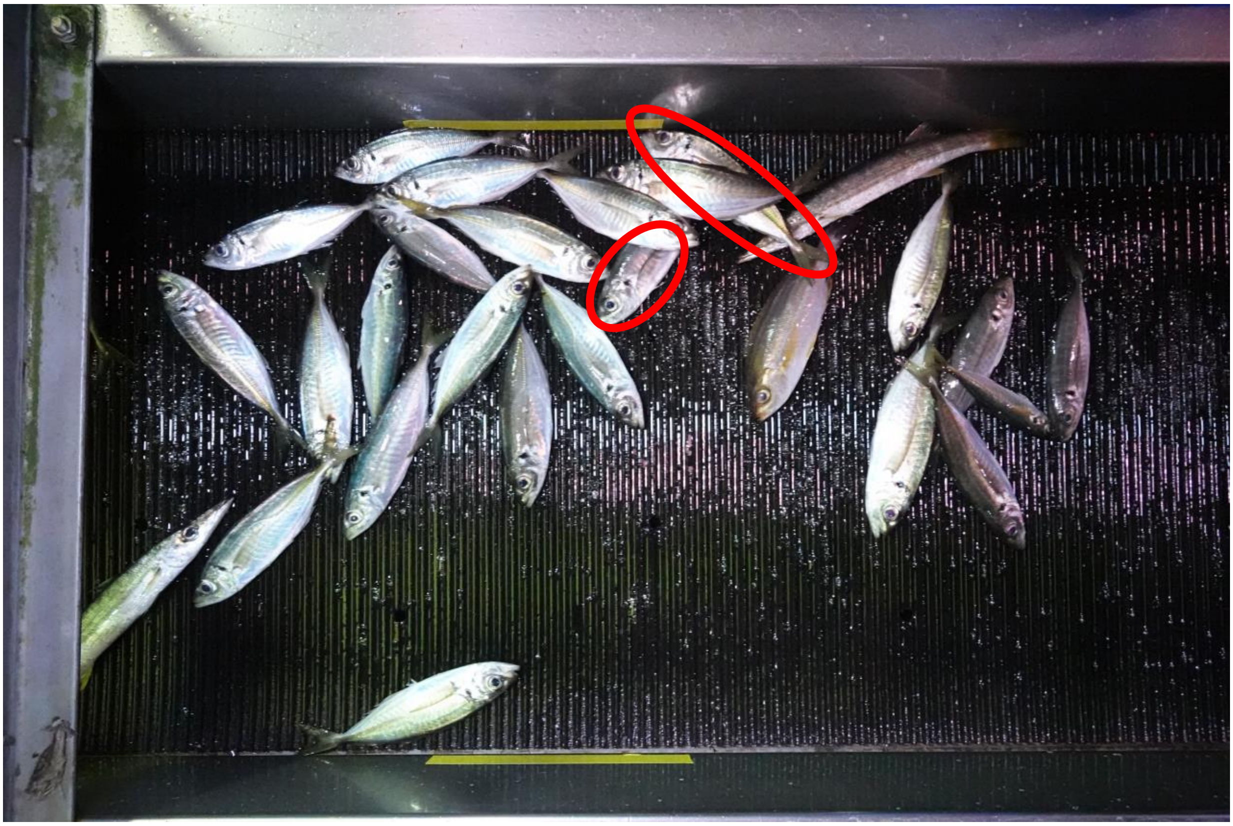

Supplemental Figure 3. Example of Japanese multiple species classes. Classes not evaluated due to lack of test data were excluded. (b) "Tobiuo“ class. Either of the following species; Exocoetidae. (c) "Aji “ class. Either of the following species; *Alectis ciliaris*, *Atropus hedlandensis*, *Caranx papuensis*, *Caranx sexfasciatus*, *Decapterus akaadsi*, *Decapterus macarellus*, *Decapterus macrosoma*, *Decapterus maruadsi*, *Decapterus muroadsi*, *Decapterus tabl*, *Decapterus* spp., *Selar crumenophthalmus*, *Trachurus japonicus*, *Uraspis helvola*.

(d)

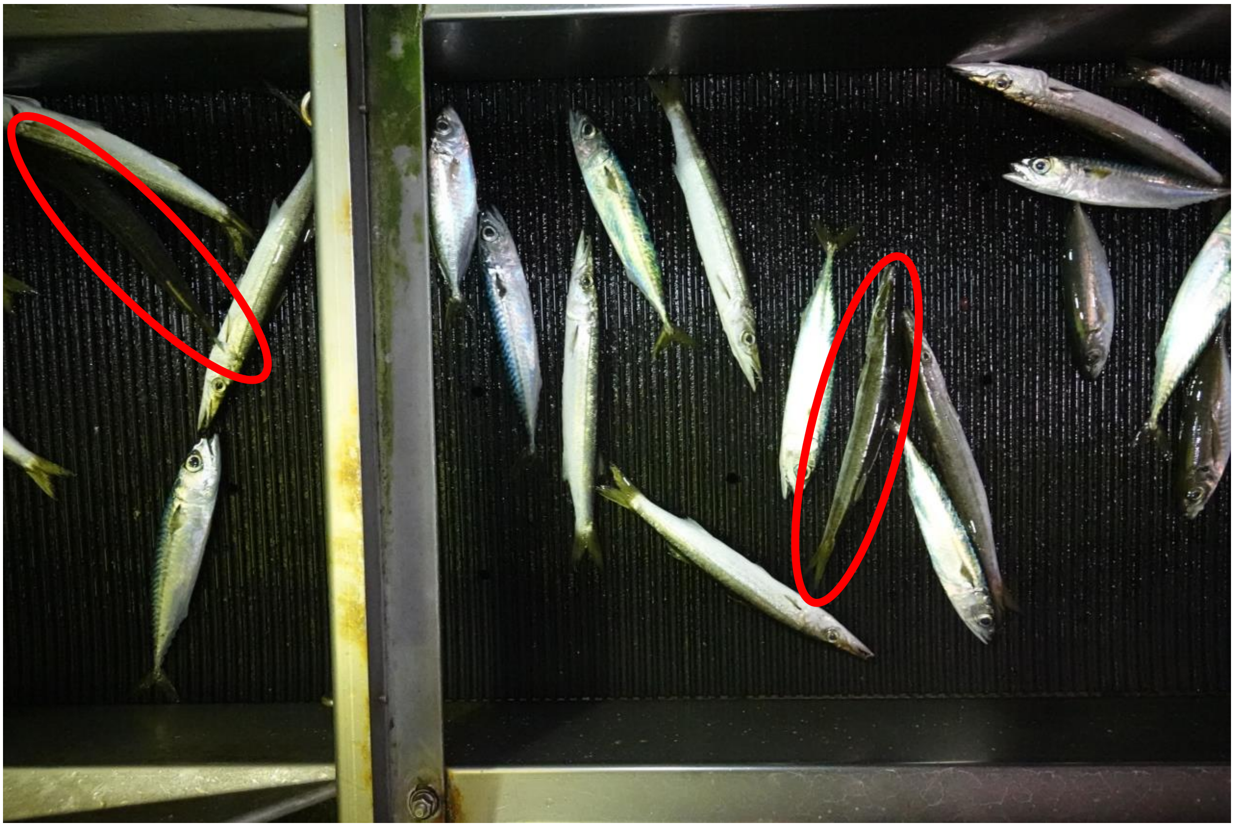

(e-1)

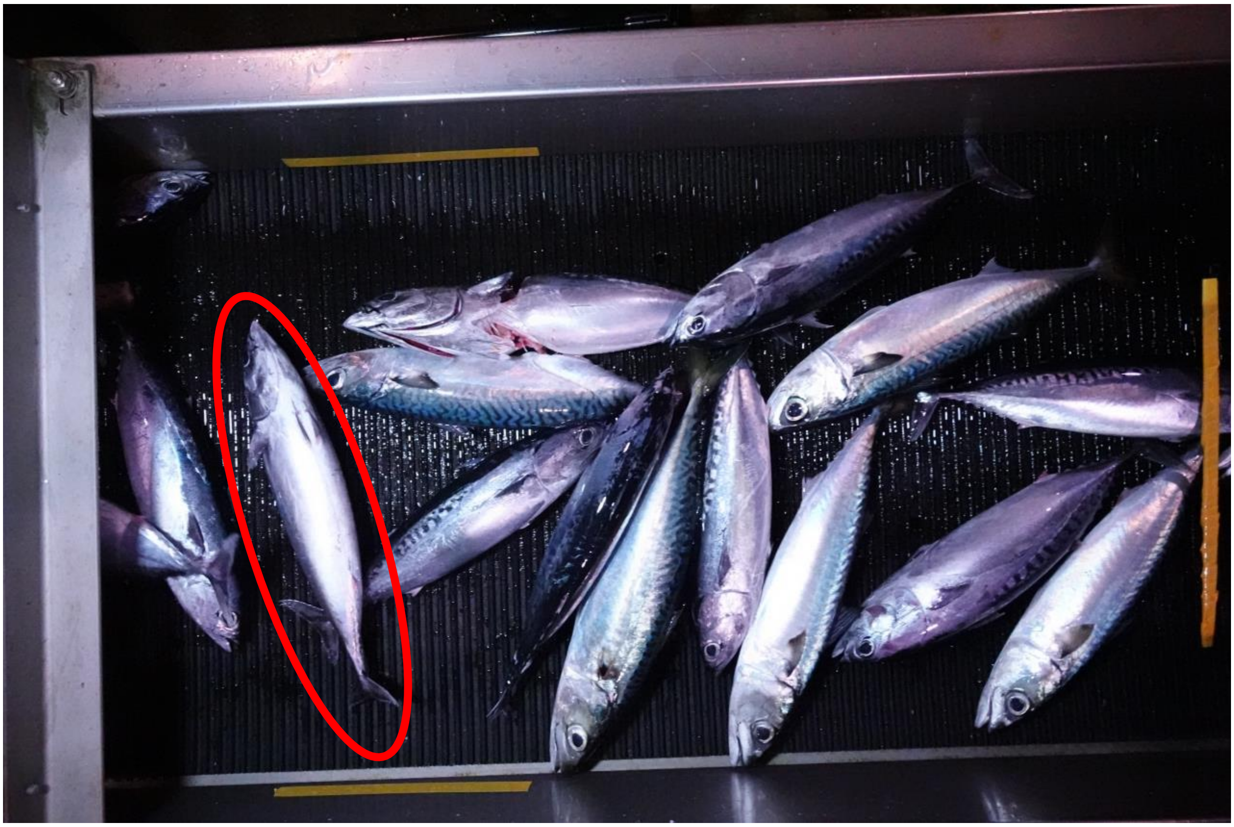

Supplemental Figure 3. Example of Japanese multiple species classes. Classes not evaluated due to lack of test data were excluded. (d) “Kamasu“ class. Either of the following species; *Sphyraena pinguis*, *Sphyraena* sp., *Promethichthys Prometheus*. (e-1) “Soudagatsuo“ class. Either of the following species; *Auxis rochei rochei*, *Auxis thazard thazard*.

(e-2)

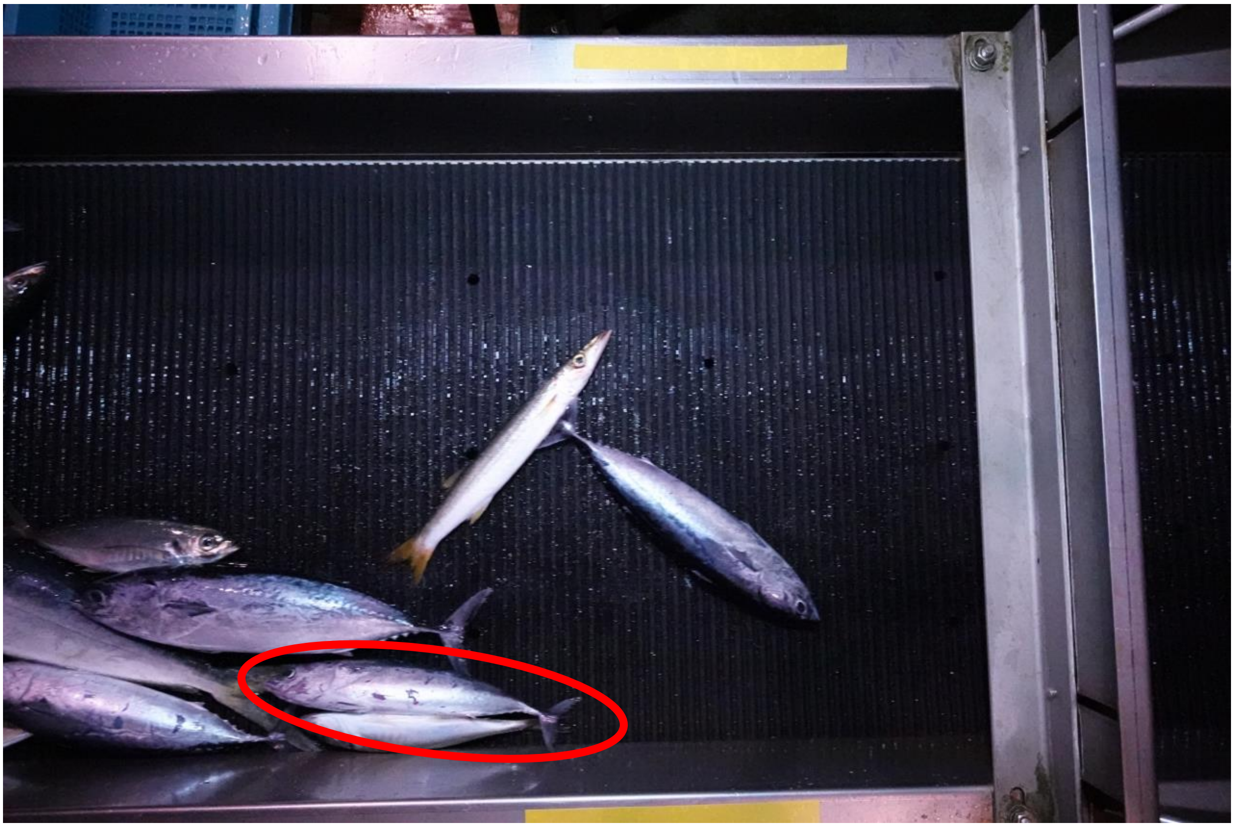

(f)

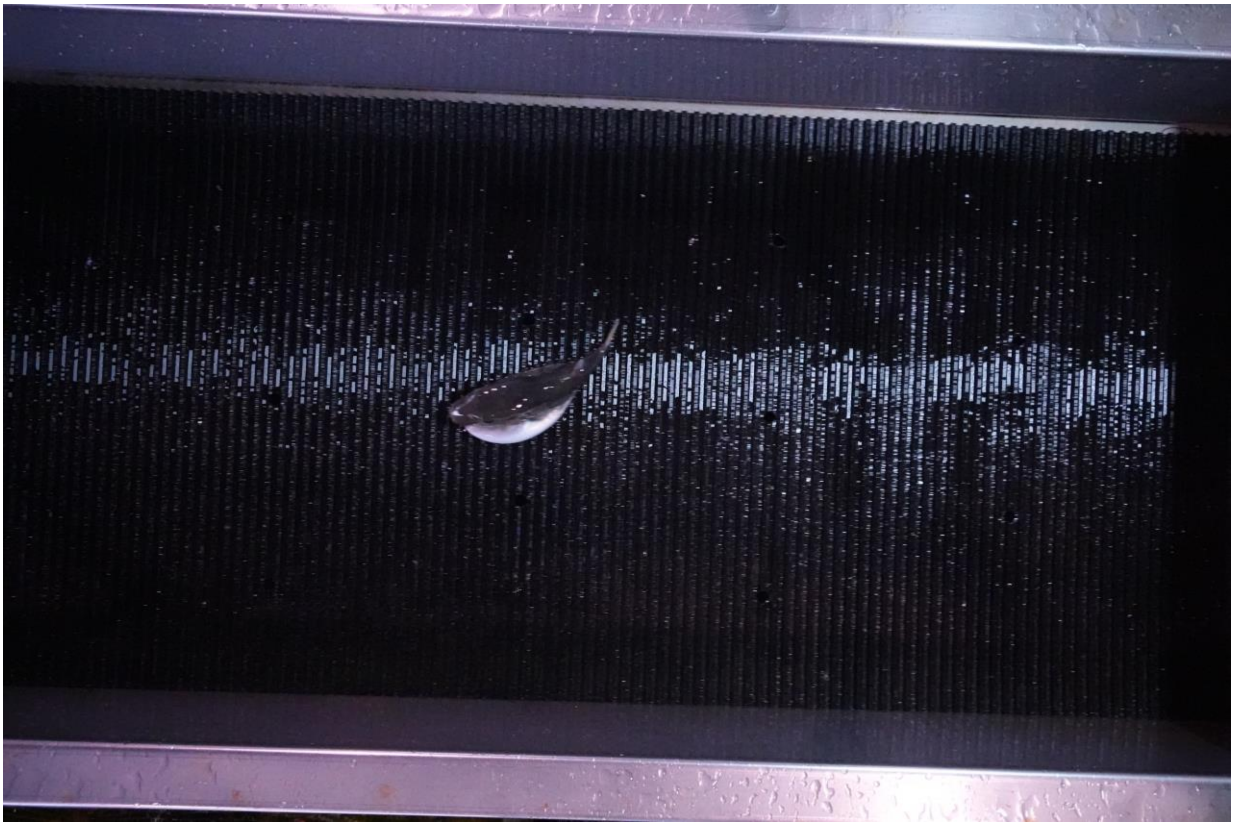

Supplemental Figure 3. Example of Japanese multiple species classes. Classes not evaluated due to lack of test data were excluded. (e-2) “Soudagatsuo“ class. Either of the following species; *Auxis rochei rochei*, *Auxis thazard thazard*. (f) “Fugu “ class. Either of the following species; Tetraodontidae.
